# A genetic toolkit to reduce wheat immunogenicity and incidence of celiac disease

**DOI:** 10.64898/2026.06.23.734071

**Authors:** Maria G. Rottersman, Debbie Laudencia-Chingcuanco, Wenjun Zhang, Maria Helena Guzman-Lopez, Jeanie W. Lin, Junli Zhang, Celine Caseys, German Burguener, Sewon Kim, Xiaoqin Zhang, Ural Yunusbaev, Eduard Akhunov, Jong-Yeol Lee, Jorge Dubcovsky

**Author notes:** Co-corresponding authors: Jorge Dubcovsky.; Debbie Laudencia-Chingcuanco.; Jong-Yeol Lee. The first three authors contributed equally to this work.

## Abstract

Celiac disease (CeD) is an immune-mediated condition triggered by wheat gluten in genetically predisposed individuals. The immune reaction in people with CeD is driven by particular gluten amino acid sequences, or immunogenic epitopes. Some of these epitopes elicit strong immune responses in the majority of CeD patients and are designated as immunodominant epitopes. Previous research has shown correlations between the amount of immunogenic wheat epitopes consumed and the onset of CeD, suggesting that reducing wheat immunogenic epitopes may reduce CeD incidence at the population level. Gluten consists of gliadins and glutenins, with gliadins having the majority of the immunodominant epitopes and glutenins playing a major role in dough strength and breadmaking quality (BMQ). This study used radiation-induced deletions, chemical mutagenesis, and natural variation in wheat (*Triticum aestivum*) to generate genetic stocks with reduced immunogenic epitope content. Most lines were developed in the wheat cultivar Summit, for which we produced a full genome assembly and annotation. We used exome capture to characterize these deletions and identify prolamins located within and outside the deletions. We combined different deletions and developed molecular markers to facilitate their deployment. For chromosome arms 1BS and 1DS, we generated two alternative lines: one lacking immunogenic epitopes for the development of CeD-safe genetic stocks for research purposes, and another retaining selected glutenins for breeding commercial lines with reduced immunogenicity and adequate BMQ. By making these non-transgenic genetic stocks publicly available, we aim to accelerate the development of wheat varieties with reduced immunogenicity and, eventually, a fully CeD-safe wheat.

## 1. Introduction

Wheat is one of the earliest domesticated crops (1, 2) and remains a major global source of calories and protein. The primary storage proteins in wheat grains are called prolamins or gluten proteins and are subdivided into two major functional groups: glutenins and gliadins. Together, these proteins form the viscoelastic gluten network responsible for the unique rheological properties of wheat dough.

Glutenins form the backbone of the gluten polymer and are classified into high- and low-molecular-weight glutenin subunits (HMW-GS and LMW-GS). These proteins are linked by interchain disulfide bonds formed through cysteine residues and are essential for dough elasticity and strength (3, 4). HMW-GS are encoded by x- and y-type genes at the *GLU1* loci on the long arms of chromosomes 1A, 1B, and 1D (5). LMW-GS are encoded by multigene families clustered at the *GLU3* loci on the short arms of the same chromosomes. In some cultivars, an additional LMW-GS gene occurs at the *GLU-B2* locus proximal to *GLU-B3* (6, 7).

Gliadins contribute primarily to dough extensibility (8) and are classified into α-, γ-, δ-, and ω-gliadins, based mainly on their electrophoretic mobility (4, 9). The α-gliadin genes are clustered at the *GLI2* locus on chromosome arms 6AS, 6BS and 6DS (10), whereas the γ-, δ-, and ω-gliadins are encoded at the *GLI1* loci on chromosome arms 1AS, 1BS and 1DS. Because of tight linkage, *GLI1* and *GLU3* are often considered a combined *GLI1/GLU3* locus. Two minor loci, *GLI-A3* and *GLI-B3*, are also located on chromosome arms 1AS and 1BS, 10-20 Mb proximal to *GLI1/GLU3*. These loci generally contain one or two functional ω1,2-gliadin genes combined with variable numbers of pseudogenes, including LMW-GS pseudogenes. In the few lines where these LMW-GS are functional, the locus has been designated as *GLU-B2* (6, 7).

Most ω-gliadins lack cysteine residues, while α-gliadins typically contain six and δ- and γ-gliadins contain eight. These cysteines form intramolecular disulfide bonds, preventing gliadins from forming covalent bonds with the gluten polymer. Instead, gliadins interact with the gluten polymer via hydrogen, ionic, and/or hydrophobic bonds (4). However, non-canonical gliadins with an odd number of cysteines can form disulfide bonds with the gluten polymer, acting as chain terminators and reducing polymer size and gluten strength (11–14). Deletion of all α-gliadins at the *GLI-D2* locus, including variants with seven cysteines, have improved gluten strength and breadmaking quality (BMQ) (12, 13). This result indicates that some immunogenic epitopes can be eliminated without compromising quality and, in some cases, even improving it. Nevertheless, excessive gliadin elimination may impair extensibility.

Despite their favorable effects on pasta and breadmaking quality, gluten proteins can trigger adverse immune reactions in susceptible individuals. Celiac disease (CeD) is a chronic autoimmune disorder affecting approximately 1% of the population (15). Genetic predisposition to CeD is strongly associated with the human leukocyte antigen alleles *HLA-DQ2* and *HLA-DQ8* (16), which occur in 30-40% of individuals (17). At present, strict adherence to a gluten-free diet remains the only effective treatment, which is challenging because of the widespread presence of wheat gluten in foods (18).

To reduce CeD risk, wheat geneticists have explored multiple strategies to eliminate immunogenic gluten components. Early studies used Chinese Spring (CS) large chromosomal deletions encompassing the *GLI1/GLU3* or *GLI2* loci (19) to evaluate their effects on immunogenicity and end-use quality (20). Smaller deletions including *GLI-B1*(21), *GLI-D1* (22), *GLI-A2*, and *GLI-D2* loci (23) were created to eliminate specific gliadins. More recently, RNA interference and genome-editing approaches, including CRISPR/Cas9, have enabled targeted modifications of various gliadin classes (24–29).

Although these studies have generated valuable information and useful genetic resources, their impact has been limited by their fragmented nature and, in some cases, restricted access to genetic materials. In this study, we developed a publicly available set of genetic stocks carrying deletions and mutations across all major and minor prolamin loci. Most of these resources were generated in the spring wheat cultivar Summit, for which we generated a high-quality genome assembly, curated prolamin annotations, and RNA-seq expression profiles during grain development.

The wheat lines developed in this study were designed to address two primary objectives. The first objective is the development of genetic stocks lacking immunogenic epitopes at each of the prolamin loci. Stacking these CeD-safe loci can be used to generate genetic stocks without immunogenic epitopes for evaluating the immunogenicity and functionality of introgressed prolamin loci and/or transgenic synthetic glutenins with edited epitopes.

The second objective, hereafter referred to as the breeding objective, is to develop wheat lines with reduced immunogenicity while maintaining acceptable end-use performance. Although these lines will not be fully safe for CeD patients, they could reduce the risks associated with inadvertent gluten exposure and potentially contribute to lowering CeD incidence.

Epidemiological studies have shown that immunogenic epitope intake correlates with CeD incidence (30, 31) and that a reduced exposure may delay disease onset (32). These findings suggest that a breeding strategy aimed at reducing the abundance of immunogenic epitopes, particularly immunodominant epitopes, could help decrease CeD incidence at the population level.

## 2. Methods

### 2.1. Sequencing, assembly, and prolamin annotation of the Summit genome

The Summit genome was sequenced to provide a reference for the characterization of the induced deletions. Summit (Express//Tadorna/Proband 775) is a commercial hard-red spring wheat cultivar (expired PVP 200200239) with a high yield potential and good BMQ. DNA extraction, quantification, and genomic library construction were performed as described previously (33). PacBio HiFi reads (∼398 Gb, 27.3X) were assembled *de novo* using Hifiasm v0.25.0 with default parameters (34). Contigs were scaffolded using Ragtag (35) with the ‘Kariega’ cultivar genome (36) as a reference. HiFi reads were remapped to the assembly using minimap2 to assess coverage in 1 Mb windows. Telomeric and centromeric motifs were identified using seqkit (37). Assembly completeness was evaluated with BUSCO v6.00 (38) using the poales_odb10 dataset. Potential contamination was assessed using the NCBI Foreign Contamination Screen toolkit (FCS) (39).

High-confidence (HC) gene models from CS genome version 2.1 (CS2.1) (40) were mapped to the Summit genome assembly using Liftoff with the “copies” parameter (41). Annotation was focused on HC genes with a “valid_ORF” tag. Duplicated genes were identified using the “extra_copy_number” tag.

Summit prolamin genes were manually curated using BLASTN searches with known CS and Fielder sequences, identification of conserved start/end regions, translation analysis, and characterization of premature stop codons, and frameshifts. Gliadins were classified according to cysteine number, diagnostic motifs, and repetitive motifs. Sequential Summit IDs were assigned within each chromosome and class.

### 2.2. RNA isolation and RNA-seq analysis

Summit plants were grown in a greenhouse under natural light supplemented with LEDs to maintain a 16 h photoperiod. One spike from each plant was tagged at anthesis and developing grains were collected at 14, 21, 28, and 35 days post-anthesis (dpa) and immediately frozen in liquid nitrogen. RNA was isolated using TRIZOL (Invitrogen, Carlsbad, CA), treated with RQ1 DNAse (Promega, Madison, WI), filtered using RNAeasy (QIAGEN Inc, Germantown, MD), and quality assessed using Agilent 2100 Bioanalyzer (Agilent, Santa Clara, CA). Sequencing was performed by Novogene (Sacramento, CA).

RNA-seq was performed for two biological replications per time point on an Illumina platform, generating an average of 42 million paired-end 150-bp reads per sample, with >95% above phred Q20. Reads were mapped at high stringency (0 SNP) to the curated Summit genome to estimate transcripts per million (TPM) of annotated prolamin genes. Mapping with Star v2.7.11 (42) yielded an average alignment rate of 82% and a 17-fold coverage. The relatively low alignment rate likely reflects the high mapping stringency.

### 2.3. Generation and selection of the Summit fast-neutron radiation mutants

Prolamin deletions were induced in Summit grains (M_0_) using 6 or 12 Gy fast-neutron radiation at the McClellan Nuclear Research Center (Sacramento, CA). Mutagenized kernels (M_1_) were grown in a greenhouse and grains from individual spikes were harvested separately to generate M_2_ families. M_3_ grains from individual M_2_ plants were screened for altered glutenin or gliadin profiles relative to wildtype (WT) using A-PAGE or SDS-PAGE, respectively. Deletion lines were initially characterized by RNA-seq using CS RefSeq v1.0 as reference, and lines carrying the shortest deletions at each locus were selected for exome-capture analysis.

### 2.4. Characterization of selected deletions using wheat exome capture

Selected deletion lines and WT Summit were further characterized using wheat exome capture assays (43). Sequencing reads were aligned to the Summit genome with BWA-MEM (44) at high stringency (0 SNPs). Reads mapping to multiple identical locations were split between locations and the number of reads per gene was determined.

Genes with fewer than 10 reads in WT Summit were excluded from the deletion analyses. Remaining read counts per gene were normalized by total reads per sample, and mutant/WT read ratios were calculated. Ratios near zero indicated deleted regions, whereas ratios near one indicated intact regions. Because reads from closely related homeologs may still map to deleted genes, especially within multigene families, moving averages across five neighboring genes on each side were used to smooth deletion profiles. At the deletion borders, average windows were restricted to genes inside or outside the deletion.

We also characterized *GLI-D1* (1DS) and *GLI-A2* (6AS) deletions introgressed in cv. Pegaso (23) (PI 709644). Because a Pegaso genome assembly is unavailable, exome capture reads were mapped to the Summit genome at 0-SNP stringency to minimizes alignment to incorrect homeologs or paralogs. Although this strategy removes polymorphic reads resulting in wider borders, it improves accuracy in defining deletion boundaries. To reduce distortions in highly polymorphic regions, genes outside the deletions showing large differences (>0.3) between Summit and Pegaso mapping ratios under strict (0 SNPs) versus variable SNP conditions were excluded from the deletion graphs.

### 2.5. RP-UPLC analysis

Gliadins and glutenins were prepared as described previously (45, 46). Freeze-dried gliadin and air-dried glutenin pellets were dissolved in 20% acetonitrile (ACN) containing 0.06% trifluoroacetic acid (TFA), filtered, and analyzed by RP-UPLC using an ACQUITY UPLC H-Class System (Waters Corp, Milford, MA, USA). Water containing 0.06% (v/v) TFA and ACN containing 0.06% (v/v) TFA were used as mobile-phase solvents.

Gliadins were separated using an ACQUITY UPLC® Peptide BEH C18 Column (Waters Corp, 300Å, 1.7 μm, 2.1×100 mm) with a linear binary gradient, starting at 25–33% of solvent B for 5 min, followed by 33-50% solvent B for 20 min and 50-90% solvent B for 1 min. Finally, solvent B was returned to 25% 4 min. The column temperature was 65 °C, the flow rate was 0.30 mL/min, and absorbance was measured at 210 nm. Glutenins were separated using a BEH C4 Column (Waters Corp, 300Å, 1.7 μm, 2.1×100 mm) with a linear binary gradient, starting at 25–45% of solvent B for 15 min, followed by 45-90% solvent B for 2 min. Finally, solvent B was returned to 25% 3 min. The column temperature was 65 °C, the flow rate was 0.20 mL/min, and absorbance was measured at 206 nm.

Peak integration was performed in the Empower3 system, and gliadin or glutenin abundance was expressed as the percentage of total peak area. Gliadin fractions were assigned by retention time as follows: ω5-gliadins (2–3.5 min), ω1,2-gliadins (6–7.5 min), α-gliadins (7.5–13.5 min), and γ-gliadin (13.5–22 min). HMW-GS and LMW-GS eluted at 6–8 min and 8–14 min, respectively.

All extractions were performed in triplicate, with three technical RP-UPLC replicates per extraction.

### 2.6. Polyacrylamide electrophoresis and two-dimensional gel immunoblot analyses

Gliadins and glutenins were sequentially extracted from half seeds as described previously (47). Gliadins were analyzed by A-PAGE and glutenins by SDS-PAGE (12%).

For selected lines, two-dimensional gel electrophoresis (2-DE) and immunoblotting were used to assess ω-gliadins. Total grain was extracted in 2% SDS lysis buffer following established protocols (48, 49). Protein concentrations were quantified using the Lowry assay (50). Proteins were separated by two-dimensional gel electrophoresis (2-DE) using isoelectric focusing followed by SDS-PAGE on NuPAGE 4–12% Bis-Tris gels (Invitrogen, Waltham, MA, USA), as described before (49). Gels were stained with Coomassie G-250 (Sigma-Aldrich, St. Louis, MO, USA) and digitized using a calibrated scanner. Duplicate analyses were performed for each sample.

For immunoblots, 7.5 μg of total SDS-extracted protein was separated by 2-DE and transferred onto nitrocellulose membranes as described previously (51). Membranes were hybridized with a monoclonal antibody against the ω5-peptide SRLLSPRGKELG (mono-ONT18A5) (21) that was diluted 1:5,000 and visualized using goat anti-mouse IgG, HRP conjugated (AbC-5001 AbClon) diluted 1:20,000 (22).

Membranes were also hybridized with a polyclonal antibody against ω1,2-gliadins (ARELNPSNKEL, Anti-NT1-ω) that was diluted 1:5,000 and visualized using goat anti rabbit IgG (H+L)-HRP (SA-002-500 GenDEPOT) diluted 1:20,000 (52). The membranes were washed three times with 1× PBS for 10 min each, followed by treatment with the corresponding HRP-conjugated secondary antibodies: (1) goat anti-mouse IgG (H+L) (1:10,000; AbClon, Seoul, Korea) or (2) goat anti-rabbit IgG (H+L) (1:10,000; GenDEPOT, Katy, USA). Detection of bound antibodies was revealed by a chemiluminescent substrate (West-Q Femto Clean ECL W3680-010 GenDEPOT) and a Chemiluminescence Imaging System Alliance Q9 Mircro (UVITEC, Cambridge, UK).

### 2.7. Proteomic analysis

Protein extraction and Liquid Chromatography-Mass Spectrometry (LC-MS) analyses of CS flours were performed as described before (12). Soluble prolamin fractions were analyzed against a reference including non-redundant curated prolamins from CS, Fielder and Summit together with Uniprot and CRAp contaminant databases. To evaluate expression and stability of ω1,2-gliadin proteins encoded at the *GLI3* locus, we identified LC-MS peptide hits uniquely matching the selected proteins.

## 3. Results

### 3.1. Sequencing, assembly, and annotation of the Summit genome

Because most fast-neutron deletions described in this study were generated in the wheat cultivar Summit, we produced a *de novo* Summit genome assembly to use as a reference. PacBio HiFi reads were assembled into 1,966 contigs spanning 14.60 Gb, with an N50 of 52 Mb and an average read coverage of ∼25× (Table S1A). Primary contigs were integrated into 21 pseudomolecules totaling 14.52 Gb, excluding mitochondrial DNA and 79 Mb of unplaced scaffolds grouped into an unassembled chromosome (Table S1B).

We recovered 4,894 of the 4,896 Benchmarking Universal Single-Copy Orthologs (BUSCO v6) for the order Poales (99.96% completeness, Table S1C), indicating a highly complete assembly. Telomeric repeats (>10 tandem TTTAGGG motifs) were identified at both ends of 17 chromosomes and at one end of the remaining four chromosomes. All 21 chromosomes contained centromeric regions identified using previously described wheat centromeric motifs (53). Alignment of the Summit assembly with the cv. Kariega genome (36) revealed strong collinearity (Fig. S1), further supporting assembly quality.

Gene annotation was transferred from CS version 2.1 high-confidence gene set (40), resulting in 110,826 aligned genes, including 99,579 valid ORFs (Table S1D). New consecutive Summit gene IDs were assigned for each pseudomolecule. Manual curation identified 78 complete prolamin genes, encoding 5 HMW-GS (*GLU-A1a*: Ax1, *GLU-B1i*: Bx17 and By18, *GLU-D1d*: Dx5 and Dy10), 13 LMW-GS, 11 γ-gliadins, 2 δ-gliadins, 12 ω-gliadins, and 35 α-gliadins (Table S2). In addition, 67 putative prolamin pseudogenes were manually annotated. Summit genomic data is available at NCBI bio-project PRJNA1438777. The numbers of known CeD immunogenic epitopes (54, 55) present in the prolamin proteins are also listed in Table S2.

To estimate prolamin gene expression, we performed RNA-seq on Summit developing grains collected at 14, 21, 28, and 35 dpa (Table S3). Transcript abundances were highly correlated between biological replicates at all time points (Table S3), so average expression values are reported in Table S2. Expression profiles across developmental stages are summarized in Fig. S2 and Table S2. Peak expression for α-gliadins, δ-gliadins, ω5-gliadins, HMW-GS, and some γ-gliadins and LMW-GS occurred at 21 dpa, whereas most γ-gliadins and LMW-GS peaked earlier (around 14 dpa). In contrast, ω1,2-gliadins showed later peak expression between 21 and 28 dpa (Fig. S2A). Similar temporal expression patterns between γ-gliadins and LMW-GS were also reflected in a Principal Component Analysis (Fig. S2B).

In summary, the Summit genome assembly, curated prolamin annotations, and accompanying RNA-seq dataset provide valuable resources for characterization of the deletion lines and expand genomic resources available for a productive spring wheat cultivar adapted to fall planting in the Mediterranean climate of California’s Central Valley.

### 3.2. Nomenclature for mutants and deletion lines

The deletion lines and natural prolamin alleles characterized in this study are summarized in Table 1. Deletion lines are designated by the chromosome arms where they are located with superscript letters indicating origin (S = Summit, K or KRO = Kronos, PEG= Pegaso, RIL= RIL143). Deletion sizes were estimated by exome capture and alignment to the Summit genome (Table S4). Mutations in individual prolamins are indicated by the locus name, with the “null” superscripts denoting non-functional alleles. Locus names are indicated in italic, whereas line names are not. Lines with combinations of all homeologs for one locus are designated by the locus name with no genome identification and a “null” superscript (e.g. the line with no HMW-GS is designated glu1^null^). Combinations of different loci are designated using the acronym CPD for “Combined Prolamin Deletion” with sequential numbers as described in Table 1.

**Table 1.**
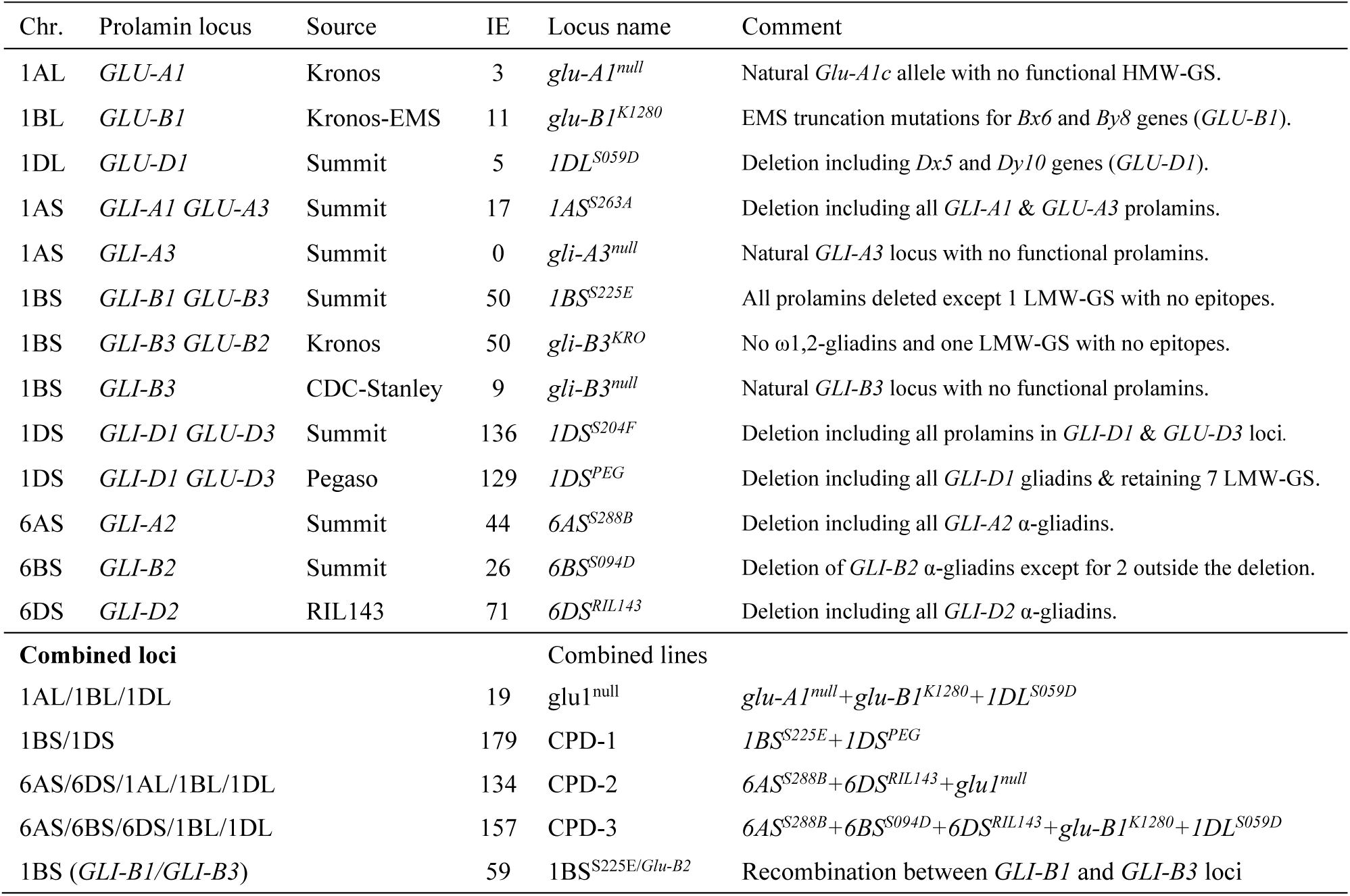
Genetic stocks generated in this study describing the affected prolamin loci, the number of immunogenic epitopes (IE) eliminated (based on Summit sequence, total IE 372), and the presence of residual prolamins.

### 3.3. The *GLU1* loci: Generation of a Summit line without HMW-GS

This section focuses on the *GLU1* loci and the development of a Summit line lacking functional HMW-GS. Among the Summit fast-neutron lines with deletions encompassing both *Dx5* and *Dy10* genes, we selected the short deletion *1DL*^S059D^. Comparison of normalized exome-capture read ratios between 1DL^S059D^ and Summit WT indicated that the deletion spans 6.60–6.85 Mb, extending from 413.21–413.42 Mb to 420.02–420.06 Mb on chromosome arm 1DL (Fig. 1A, Table S5). Deletion borders are provided as ranges because they were estimated from flanking genes coordinates. The *1DL*^S059D^ deletion removed *Dx5*, *Dy10*, and 84 additional non-prolamin genes (Table S5).

**Fig. 1.**
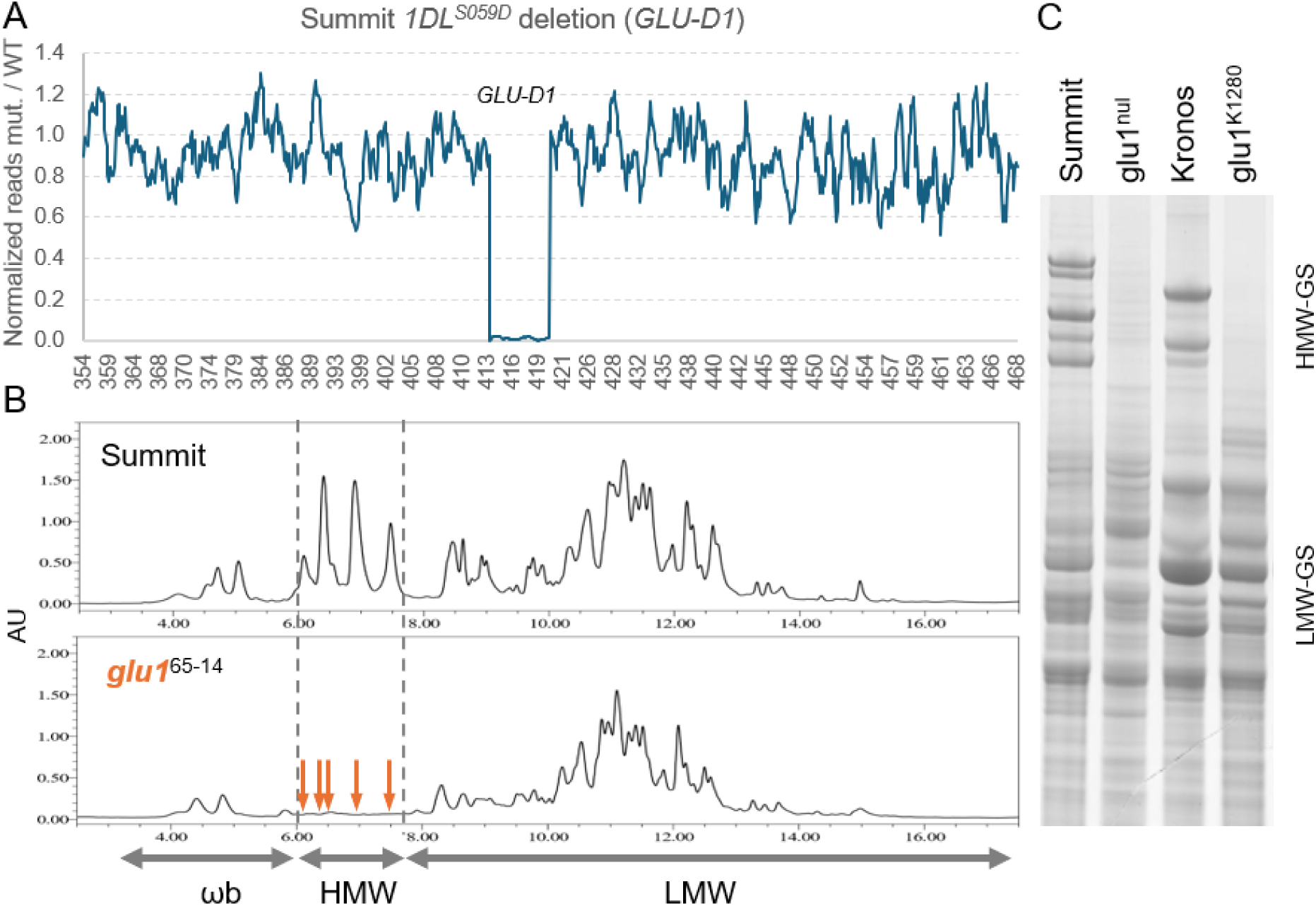
Characterization of *GLU1* deletions. (**A**) Ratio between exome capture normalized number of reads per gene in 1DL^S059D^ and Summit WT, mapped to the Summit genome at high stringency (0 SNPs). Values in the x-axis indicate positions in Mb along chromosome 1D. (**B**) RP-UPLC analysis of glutenins for Summit WT and glu1^null^ lacking all functional HMW-GS. The x-axis indicates elution times (min) and the y-axis the intensity of the signal in absorbance unit (AU). Arrows indicate missing peaks. HMW= high molecular weight glutenins, LMW= low molecular weight glutenins, and ωb= omega bound gliadins (likely ω-gliadins with one cysteine that facilitates covalent bonding to the gluten polymer). (**C**) SDS-PAGE analysis of glutenins for Summit, Kronos, and the deletion lines glu-B1^K1280^ and glu1^null^, both lacking all functional HMW-GS.

Non-functional *GLU-A1* and *GLU-B1* alleles were introgressed into 1DL^S059D^ from the tetraploid wheat (*Triticum turgidum* ssp. *durum* cv. Kronos) EMS mutant line glu-B1^K1280^ (PI 692252). This line carries premature stop codons in Bx6 (Q117*) and By8 (R64*), truncating >85% of the encoded proteins (56), together with the natural *Glu-A1c* allele (hereafter *gli-A1*^null^), which is common in tetraploid wheat (57). This locus includes an Ax pseudogene with a premature stop codon (Q406*) and an Ay pseudogene disrupted by a WIS retrotransposon insertion. The *glu-A1^null^* and *glu-B1^K1280^*loci were introgressed into the hexaploid line 1DL^S059D^ by four backcrosses (Fig. S3), generating line glu1^null^ lacking functional HMW-GS genes (Fig. 1B). Molecular markers used to introgress and combine the three non-functional *GLU1* loci are listed in Table S6.

### 3.4. The *GLI2* loci: Generation of a Summit line depleted of *α*-gliadins

The *GLI2* loci in Summit contain 13 complete α-gliadins in *GLI-A2*, 13 in *GLI-B2* and 9 in *GLI-D2* (Fig. 2). The Summit chromosome arm 6AS deletion, designated *6AS^S288B^*, extends from 25.72–25.98 Mb to 29.18–29.22 Mb, spanning 3.20–3.50 Mb. The *6AS^S288B^*deletion includes all 13 α-gliadin genes (44 immunogenic epitopes), 22 of the 23 α-gliadin pseudogenes, and six non-prolamin genes (Table S7, Fig. 2A-B). We also characterized a previously described *GLI-A2* deletion introgressed into cv. Pegaso from cv. Reader (23), designated *6AS^PEG^*. Exome capture data showed that *6AS^PEG^* is substantially larger (41.56–41.68 Mb) than *6AS^S288B^* and encompasses all 6AS α-gliadin genes plus 594 non-prolamin genes (Fig. S4, Table S7), so it was excluded from subsequent studies.

**Fig. 2.**
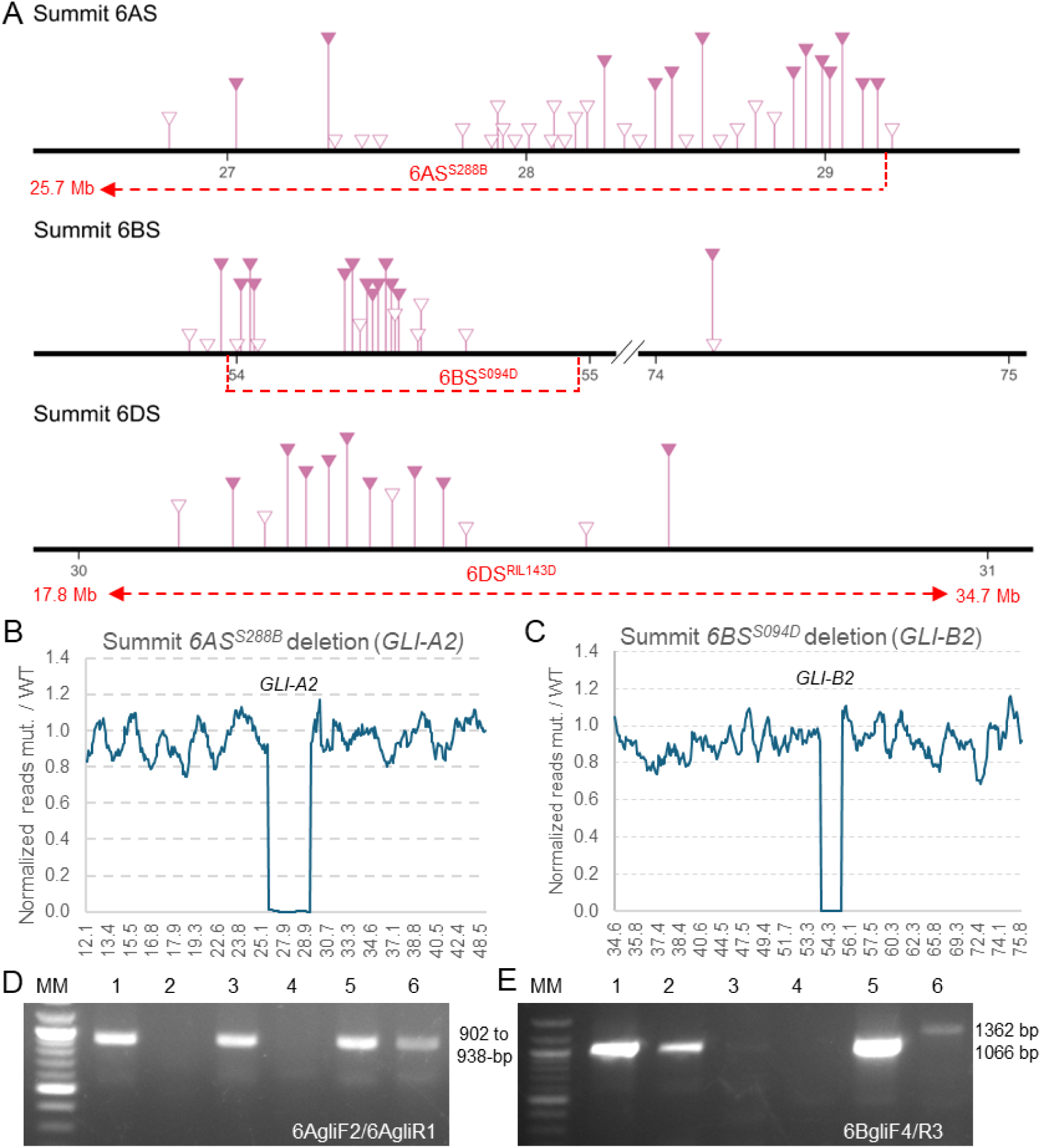
Characterization of *GLI2* deletions. (**A**) Distribution of Summit α-gliadin genes (dark pink triangles) and pseudogenes (empty triangles) on the *GLI-A2, GLI-B2* and *GLI-D2* loci. Numbers indicate coordinates in Mb based on the Summit genome. Dotted red lines indicate deletions *6AS^S288B^* and *6BS^S094D^* in Summit, and *6DS^RIL143^*(described previously (12)). (**B** and **C**) Ratios between exome capture normalized reads of mutants over WT along chromosomes 6A and 6B for deletions *6AS^S288B^*(**B**) and *6BS^S094D^* (**C**). Values in the x-axis indicate chromosome positions in Mb. (**D** and **E**) Markers 6AgliF2/R1 (**D**) and 6BgliF4/R3 (**E**) Lines: 1. Summit WT, 2. 6AS^S288B^, 3. 6BS^S094D^, 4. CPD-3 (combined *6AS^S288B^+ 6BS^S094D^+ 6DS^RIL143^ + glu-B1^K128^ + 1DL^S059D^* deletions), 5. glu1^null^, 6. Kronos.

To facilitate the introgression of *6AS^S288B^*, we developed primers 6AgliF2/R1 (Table S6), which amplify multiple *Gli-A2* α-gliadins as fragments of similar size (902–938 bp depending on genotype). No amplification was detected in plants homozygous for the deletion (Fig. 2D). We also developed the linked codominant marker 6AufF1/R1(Table S6), located 1.15–1.20 Mb proximal to the deletion, to facilitate selection of heterozygous introgression lines (Table S6).

The *6BS^S094D^* deletion is comparatively small (0.72–0.87 Mb), extending from 53.96–53.98 to 54.70–54.82 Mb (Table S7, Fig. 2A and C). This deletion can be traced with dominant marker 6BgliF4/R3, developed for one of the deleted α-gliadin genes (Table S6). This marker amplifies a 1,066-bp fragment in hexaploid wheat and of 1,362-bp fragment in tetraploid wheat, both absent in plants homozygous for the *6BS^S094D^* deletion (Fig. 2E).

The *6BS^S094D^* deletion removes 11 α-gliadin genes (26 immunogenic epitopes), 7 pseudogenes, and four non-prolamin genes (Table S7 and Fig. 2A). However, two α-gliadins remain outside the *GLI-B2* deleted region: *αGLI-B2.1* and *αGLI-B2.13* (Table S2). RNA-seq analyses showed that both genes are expressed in the developing grain (Table S2) and encode proteins containing immunogenic epitopes. The proximal gene, *α-GLI-B2.13* (6BS: 74.16 Mb), is ∼19 Mb from the deletion (Table S7) and could potentially be replaced by recombination with a natural pseudogene (Q101*) identified in Cadenza, Claire, Julius, Attraktion, and SY-Mattis. By contrast, *α-GLI-B2.1* is tightly linked to the deletion (< 21 Kb, Table S7), making elimination by recombination unlikely. Thus, development of a CeD-safe *6BS^S094D^* locus will require editing of the α-gliadin genes outside the deleted region.

We also remapped the exome capture of *6DS^RIL143^* (12) to Summit and established that the 9 α-gliadin genes (71 immunogenic epitopes) and the 5 α-gliadin pseudogenes were all within this deletion together with 167–176 non-prolamin genes (Table S7). Based on the Summit coordinates the *6DS^RIL143^* is 14.66–16.90 Mb long and extends from 19.87–21.85 to 36.52–36.78 Mb on chromosome arm 6DS.

To generate a triple *gli2-*deletion line combined with *GLU1* knockout alleles, we first introgressed *6DS^RIL143^* (12) into glu1^null^ by four backcrosses. Dominant marker 6DgliF2/R2 and linked codominant SSR marker BARC54 (Table S6) were used to track *6DS^RIL143^* and identify a BC_4_F_2_ plant homozygous for the four non-functional alleles (*glu1^null^ 6DS^RIL143^*, Fig. S5). In parallel, glu1^null^ was crossed with 6AS^S288D^, and a plant homozygous for the four non-functional loci was selected and crossed with the line combining the *glu1^null^ 6DS^RIL143^*deletions (Fig. S5). From the F_2_ progeny, we selected a plant homozygous for *6AS^S288B^*, *6DS^RIL143^*, and *glu1^null^*, designated CPD-2 (Fig. S5). Finally, CPD-2 was crossed with 6BS^S094D^, generating line CPD-3 homozygous for all three *GLI2* deletions plus *glu-B1^K1280^* and *1DL^S059D^* alleles (*GLU-A1x* is still present) (Fig. S5). In total, 157 immunogenic epitopes were eliminated in this line.

To visualize deleted α-gliadins, we compared RP-UPLC (Fig. 3A) and A-PAGE (Fig. 3B) profiles of Summit WT and the *GLI2* deletion lines. Relative gliadin fractions were quantified from RP-UPLC data and are presented in Fig. S6 and Table S8. In Summit WT, α-gliadins represented 47.90% of total gliadins, a proportion that decreased to 40.68% in 6BS^S094D^ and to 30.26% in CPD-2, with compensatory increases in γ- and ω-gliadins (Fig. S6 and Table S8, relative %). These decreases were consistent with the RP-UPLC profiles, which lacked four peaks in 6BS^S094D^ and eight in CPD-2 (Fig. 3A). In the A-PAGE, CPD-2 lacked α-gliadin bands at the bottom of the gel, corresponding to *GLI-A2* and *GLI-D2* deletions (Fig. 3B). Line CPD-3, carrying deletions in all three *GLI2* loci, retained only two α-gliadin bands (Fig. 3B), likely corresponding to the *α-GLI-B2.1* and *α-GLI-B2.13* genes located outside the *6BS^S094D^* deletion (Fig. 2A, Table S7). Line CPD-3 was unavailable when RP-UPLC analyses were performed.

**Fig. 3.**
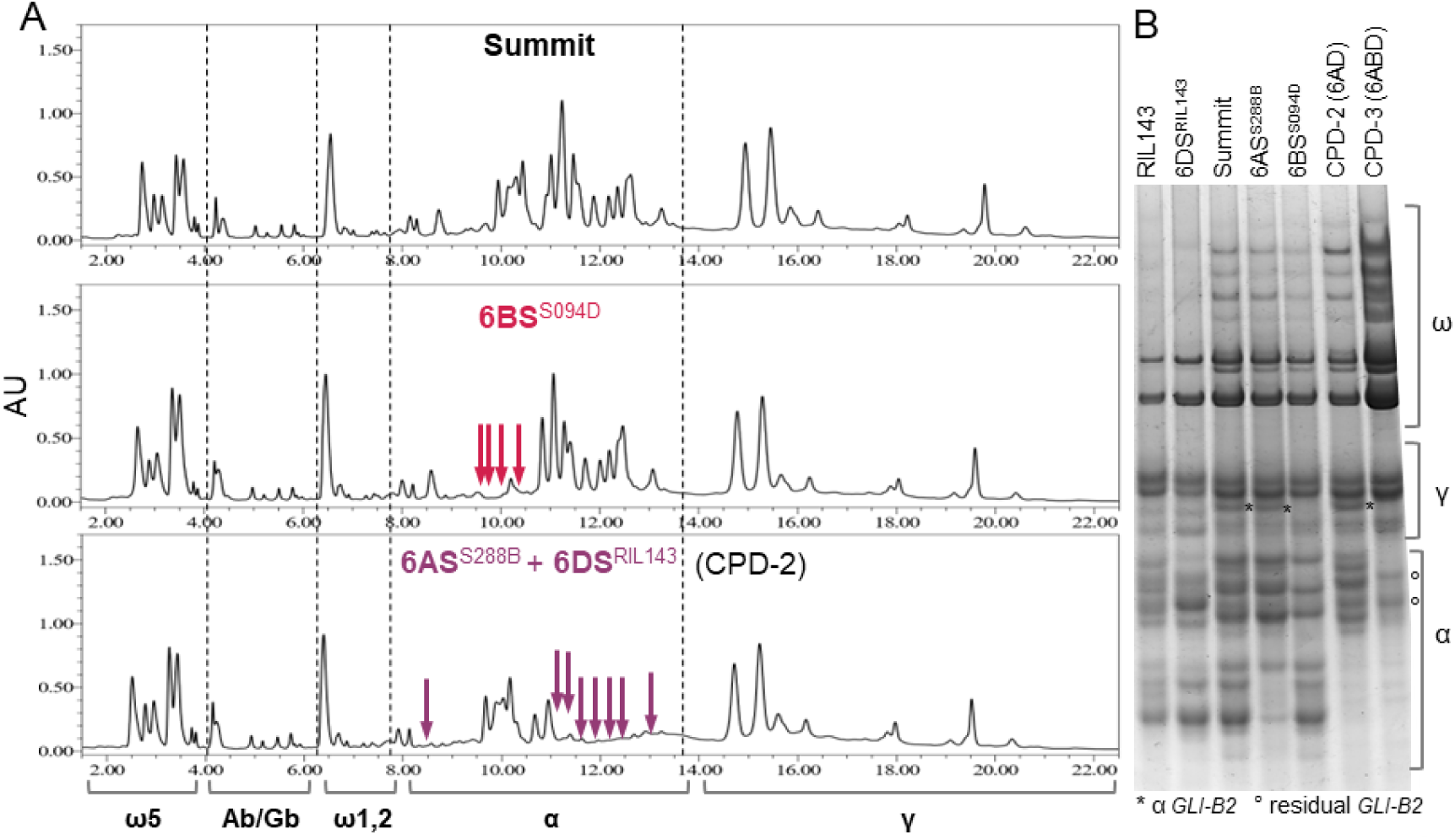
RP-HPLC and A-PAGE pattern analysis of gliadins for Summit WT and *GLI2* deletion lines. (**A**) RP-HPLC. Deleted α-gliadins in line 6BS^S094D^ are indicated by red arrows and those in CPD-2 (*6AS^S288B^ 6DS^RIL143^*) by violet arrows. (**B**) A-PAGE. The first two lanes are in RIL143 background and the last five in Summit background. Line CPD-3 combines deletions for all three *GLI2* loci. The two α-gliadin bands present in CPD-3 (marked by °) likely correspond to the two α-gliadins outside the *6BS^S094D^* deletion (Table S7). The higher intensity of the ω- and γ-gliadins in CPD-3 is the result of compensation for missing α-gliadins and not of differential loading. This line was repeated multiple times in different gels with the same result. *= indicates an α-gliadin from *GLI-B2* located in the γ-gliadin region.

### 3.5. The *GLI1/GLU3* combined loci

#### 3.5.1. GLI-A1/GLU-A3

The Summit *GLI-A1 /GLU-A3* locus includes genes encoding one δ-gliadin, three γ-gliadins, two ω5-gliadins, and three LMW-GS, which are all missing in the *1AS^S263A^* deletion (Fig. 4A, Table S2, 17 immunogenic epitopes eliminated). This deletion also includes six prolamin pseudogenes and 25 non-prolamin genes (Table S9). The *1AS^S263A^* deletion spans 1.93–2.05 Mb from 7.11–7.20 Mb to 9.13–9.15 Mb on chromosome arm 1AS (Fig. 4A-B, Table S9). To track this deletion, we developed the dominant marker 1AgliF9/R9 (Table S6), which amplifies an 877-bp fragment in plants without the deletion but no product in homozygous *1AS^S263A^*plants (Fig. 4C). A codominant CAPS marker tightly linked to *1AS^S263A^* (1AgluF3/R3 with *Hind*III digestion, Table S6) was developed to monitor the deletion in heterozygous state during introgressions.

**Fig. 4.**
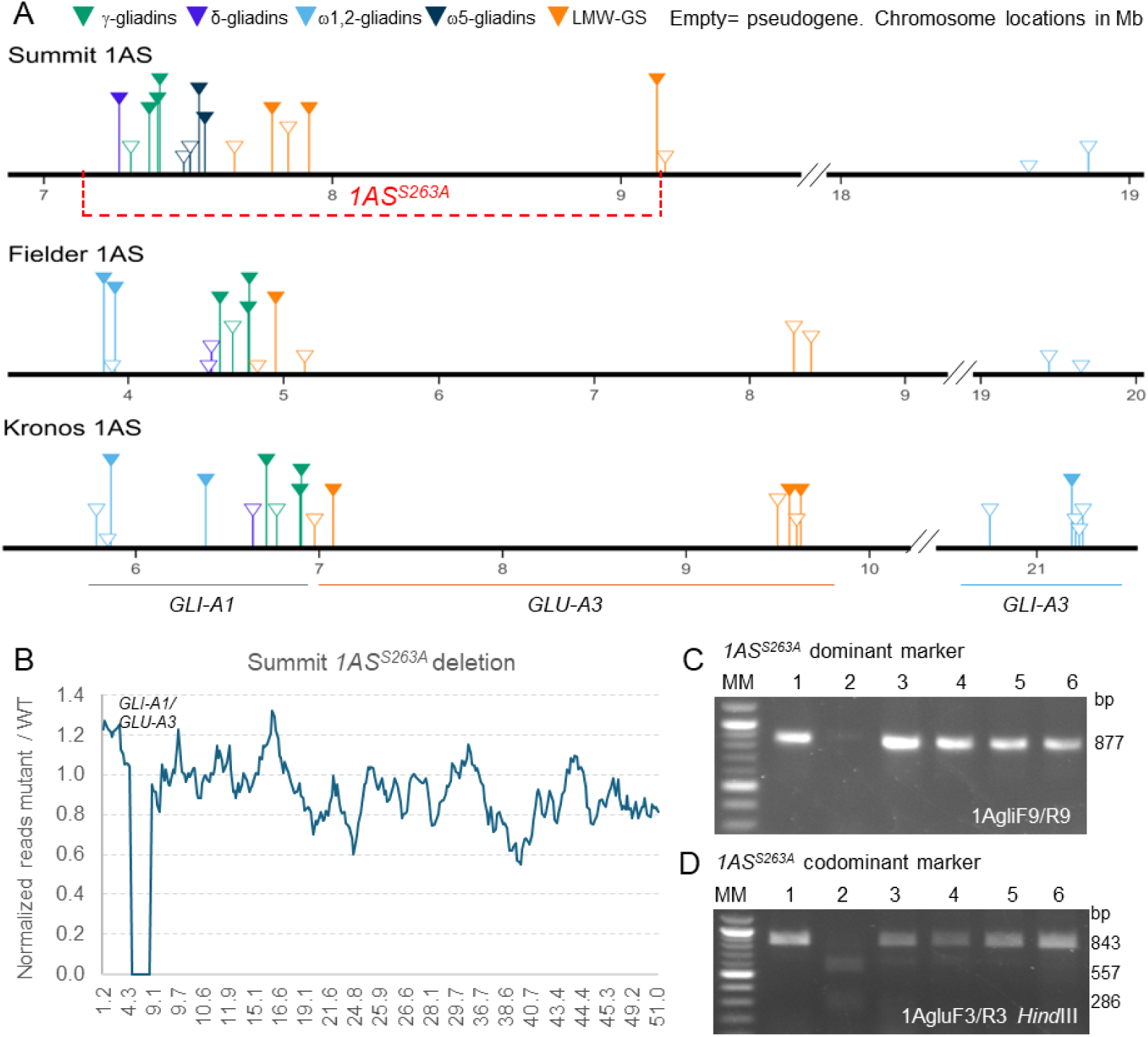
Tools for the *GLI-A1/GLU-A3* loci on chromosome arm 1AS. (**A**) Distribution of gliadins and LMW-GS on chromosome arm 1AS of hexaploid wheat cv. Summit and Fielder, and tetraploid wheat cv. Kronos. The dotted red line below Summit indicates the position of the *1AS^S263A^* deletion. Numbers indicate coordinates in Mb based on their respective genomes. There is a break in the Mb distances between the *GLI1/GLU3* and the *GLI3* loci. (**B**) Ratio between normalized read number in the exome captures from lines 1AS^S263A^ and Summit WT. Values in the x-axis indicate positions in Mb on chromosome 1A. (**C**) 1AgliF9/R9 dominant marker for the *1AS^S263A^* deletion. Lines 1. Summit, 2. 1AS^S263A^, 3. 1BS^S225E^, 4. 1DS^PEG^, 5. glu1^null^, 6. glu-B1^K1280^ (**D**) 1AgluF3/R3 – *Hind*III codominant CAPS marker for the *1AS^S263A^* deletion. Lines 1. Kronos. 2. Summit, 3. Fielder, 4. CS, 5. RIL143, 6. Pegaso.

#### 3.5.2. GLI-B1/GLU-B3

The Summit *GLI-B1/GLU-B3* locus contains genes encoding four γ-gliadins, three ω5-gliadins and four LMW-GS including 50 immunogenic epitopes (Fig. 5A). With the exception of *GLU-B3.4*, all other prolamin genes are deleted in *1BS^S225E^*. *GLU-B3.4* is highly expressed (Table S2), and the encoded LMW-GS lacks reported CeD immunogenic epitopes (54, 55).

**Fig. 5.**
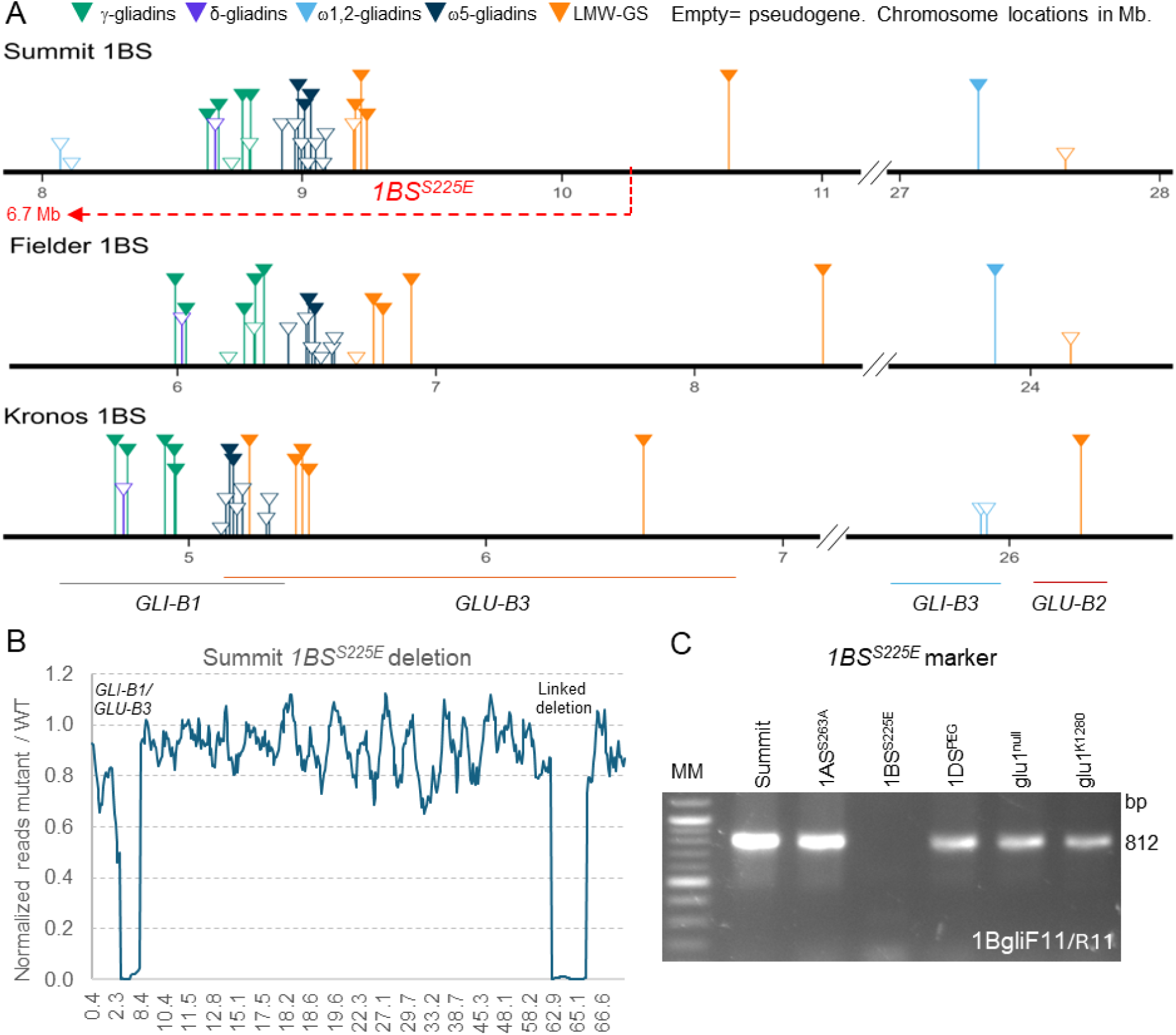
Tools for the *GLI-B1/GLU-B3* loci on chromosome arm 1BS. (**A**) Distribution of gliadins and LMW-GS on the 1BS chromosome arms of hexaploid cv. Summit and Fielder and tetraploid cv. Kronos. The dotted red line below Summit indicates the *1BS^S225E^*deletion. Numbers indicate coordinates in Mb based on their respective genomes. There is a break in the Mb distances between the *GLI1/GLU3* and the *GLI3/GLU2* loci. (**B**) Ratio between normalized read number in the exome captures of lines 1BS^S225E^ and Summit WT. Values in the x-axis indicate positions in Mb on chromosome 1B. (**C**) 1BgliF11/R11 dominant marker for the *1BS^S225E^* deletion.

The *1BS^S225E^* deletion extends from 6.74–7.16 Mb to 9.74–10.25 Mb (Fig. 5A-B), generating a 2.58–3.51 Mb deletion encompassing 44 non-prolamin genes. We developed dominant marker 1BgliF11/R11 from a γ-gliadin located within the deletion (Table S6), which amplifies an 812-bp fragment in Summit but no product in in homozygous *1BS^S225E^* plants (Fig. 5C). A secondary deletion was detected 27.3 Mb proximal to *1BS^S225E^*, between 64.92–65.69 and 69.13–69.35 Mb (Fig. 5B, Table S9). This deletion spans 3.44–4.43 Mb and includes 46 annotated non-prolamin genes (Table S9). This secondary deletion is expected to be eliminated after the *1BS^S225E^* deletion is recombined with the *GLI-B3* locus from either CDC-Stanley or Kronos (see section 3.6.2).

#### 3.5.3. GLI-D1/GLU-D3

The *GLI-D1* locus on chromosome arm 1DS is similar in Summit and Fielder, containing genes encoding four to five ω1,2-gliadins in the distal region, one δ-gliadin, four γ-gliadins, two ω5-gliadins, and six LMW-GS including 136 immunogenic epitopes (Fig. 6A). All these prolamin genes are absent in the 11.18–11.71 Mb distal deletion in Summit 1DS designated *1DS^S204F^*(Table S9, Fig. 6A-B). Besides 17 deleted prolamins and 6 pseudogenes, this deletion removed 284 non-prolamin genes (Table S9). Since *1DS^S204F^* was identified recently, it has not yet been combined with *1AS^S263A^* and *1BS^S225E^*.

**Fig. 6.**
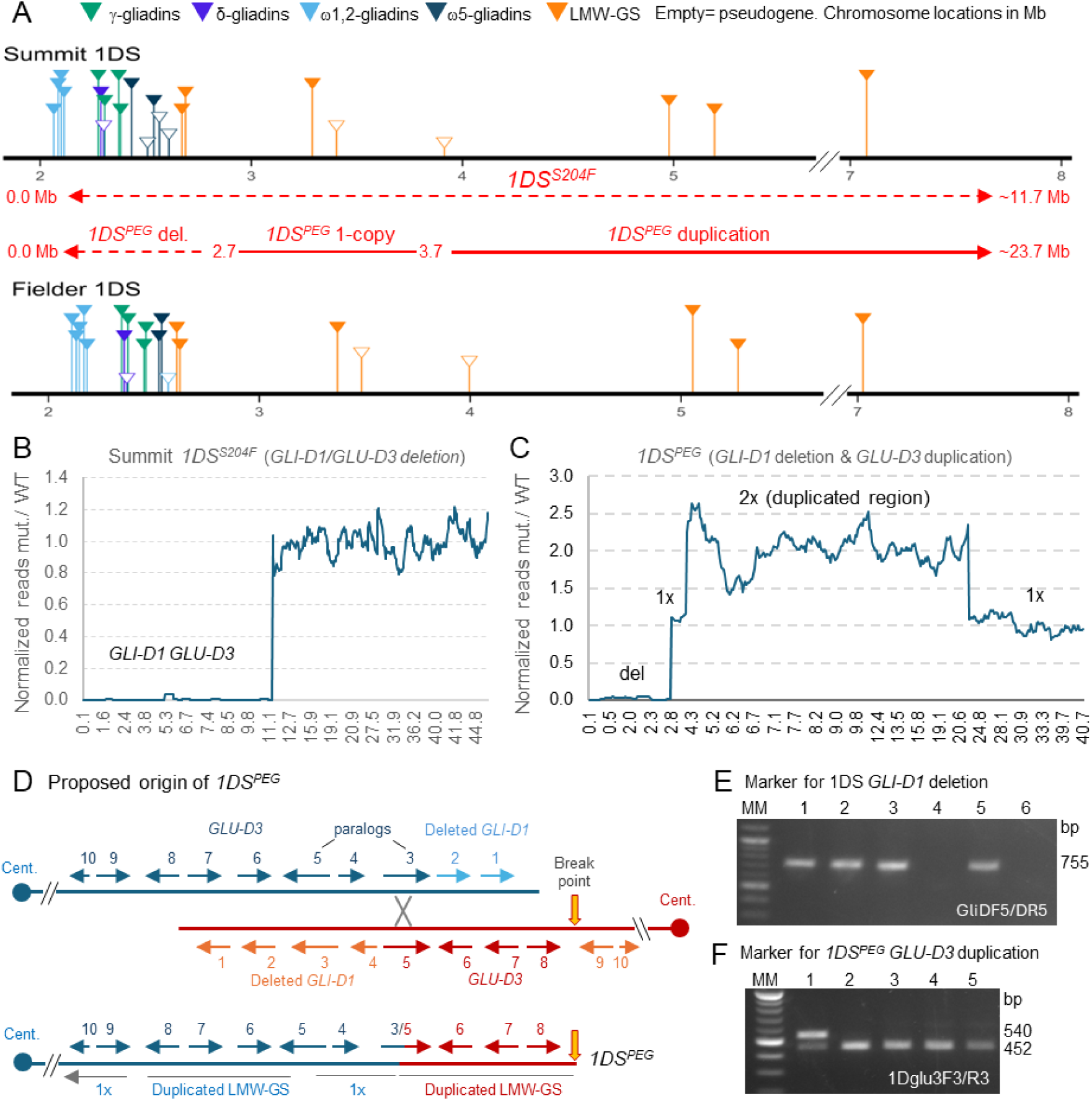
Tools for *GLI-D1/GLU/D3* prolamin loci on chromosome arm 1DS. (**A**) Distribution of gliadins and LMW-GS on chromosome arm 1DS of hexaploid cv. Summit and Fielder. Numbers indicate coordinates in Mb based on their respective genomes. The dotted red lines below Summit indicate *1DS^S204F^* deletion and the inferred deleted and duplicated regions in *1DS^PEG^*. (**B - C**) Ratios between normalized read number in exomes captures from (**B**) *1DS^S204F^* deletion including *GLI-D1-GLU-D3* and (**C**) *1DS^PEG^* rearranged chromosome. Values in the x-axis indicate positions in Mb on chromosome 1D. (**D)** Proposed mechanism for the origin of *1DS^PEG^*. Top: proposed ectopic recombination event, bottom: rearranged *1DS^PEG^*. Numbers and arrows represent hypothetical genes. (**E**) GliDF5/DR5 dominant marker for the deleted region in 1DS^PEG^. Lines 1. Summit, 2. 1AS^S263A^, 3. 1BS^S225E^, 4. 1DS^PEG^, 5. glu1^null^, 6. glu-B1^K1280^ (tetraploid, no D genome). (**F**) 1Dglu3F3/R3-*Nco*I codominant marker for the duplicated region in *1DS^PEG^*. Lines: 1. 1DS^PEG^, 2. Summit, 3. Fielder, 4. CS, 5. RIL143. Note the duplicated band in *1DS^PEG^*.

We also characterized a natural *GLI-D1* deletion discovered in the French cultivar Darius (PI 422278), which was then transferred into the Italian cultivar Pegaso (*1DS^PEG^*) together with the *6AS^PEG^* deletion described earlier (23). There is currently no Pegaso genome sequence, so exome capture reads from Pegaso were mapped to the Summit genome and distances were estimated based on Summit coordinates. Read coverage across *1DS^PEG^* revealed a complex chromosome structure consisting of a distal deletion (2.69–2.75 Mb), a single-copy region (0.90–1.07 Mb), and a duplicated segment (19.41–19.99 Mb long) that extended from 3.66–3.77 Mb to 23.17–23.65 Mb (Table S9, Fig. 6A and C). Summit 1DS gliadins and two LMW-GS were included within the deleted region (Table S9), whereas one LMW-GS mapped to the single copy region and three to the duplicated region. These results suggest the presence of 7 LMW-GS in *1DS^PEG^*.

We propose that this structural rearrangement originated from ectopic recombination between inverted paralogous genes at *GLU-D3* (Fig. 6D), generating an acentric fragment carrying *GLI-D1* (lost after meiosis) and a dicentric chromosome bridge. A bridge breakage likely produced the inverted duplicated region (Fig. 6D). Multiple heterozygous SNPs between the duplicated genes suggest an origin from divergent haplotypes.

For both *1DS^S204F^* and *1DS^PEG^* deletions, we developed a dominant marker (GligDF5/DR5del1D, Table S6) producing no PCR amplification in plants homozygous for the deletions and a 755-bp fragment in plants without these deletions (Fig. 6E). We also developed a codominant marker for a duplicated LMW-GS polymorphic for an *Nco*I restriction site (1Dglu3F3/R3-LMW-het, Table S6). *Nco*I digestion of 1DS^PEG^ PCR products produced both undigested (540 bp) and digested (452 + 88 bp) fragments, a pattern unique to 1DS^PEG^ (Fig. 6F). Using these markers, we confirmed that the rearranged *1DS^PEG^* chromosome was already present in Darius.

To assess cytogenetic stability, we analyzed 10 plants from 10 separate head-rows of a line homozygous for *1DS^PEG^*. All plants carried the *GLI-D1* deletion (Fig. S7A-B) and most of them showed the duplicated codominant marker. However, two individuals from head-row 103 were homozygous for one of the two bands (Fig. S7C), and four from head-row 108 lacked amplification entirely (Table S6, Fig. S7D). These patterns are consistent with incorrect pairing of two *1DS^PEG^*chromosomes by their inverted segments, recombination within the inverted segments (Fig. S7E), and independent breakage points of dicentric bridges. These results indicate some cytogenetic instability of *1DS^PEG^*.

To combine different *GLI1/GLU3* deletions, the 1BS^S225E^ line was crossed with an available F_1_ plant from the cross Pegaso × RIL143. An F_2_ line homozygous for both *1DS^PEG^* and *1BS^S225E^*, designated CPD-1, was selected (still segregating for the Summit, Pegaso, and RIL143 genomes). Despite the presence of two 1S deletions, this line showed normal growth and fertility.

### 3.6. The *GLI3* loci

The *GLI3* loci comprise small clusters of ω1,2-gliadines on chromosome arms 1AS and 1BS located 10–20 Mb proximal to their respective *GLI1/GLU3* loci (Table S2). A neighbor-joining tree including ω1,2-gliadins from *GLI1* and *GLI3* loci resolved separate GLI1 and GLI3 protein clusters (Fig. S8A), suggesting that *GLI3* origin predated the divergence of the A and B genomes. This result is consistent with a shared premature stop among ω1,2-gliadins from the *GLI-A3* and *GLI-B3* loci that eliminates six conserved C-terminal amino acids (Fig. S8B, Table S10).

In CS, Summit, and Fielder, the ω1,2-gliadin gene at *GLI-B3* encodes a protein with an additional premature stop codon truncating 31 C-terminal amino acids (Fig. S8B, Table S10). To determine whether the six and 31 amino acid truncations affected ω1,2-gliadin stability, we performed proteomics experiments in duplicate using CS flour and a reference wheat grain proteome (Table S11). The two experiments produced similar results and are discussed together. Combined LC-MS results identified 26 peptides (155 spectral counts) unique for GLI-A3 and 23 (107 spectral counts) unique for GLI-B3 (Table S12), representing 33.3% and 20.9% of total ω-gliadin spectral counts, respectively. These results indicate that these ω1,2-gliadin proteins are relatively abundant despite their small C-terminal truncations.

The functional ω1,2-gliadins encoded at the *GLI3* locus include multiple copies of the CeD immunodominant epitope QQPQQPFPQ (*GLI-A3*) or the related epitope LQPQQPFPQ (*GLI-B3*, Table S10), so their elimination is important for the development of CeD-safe wheats.

Fortunately, genetic maps place the *GLI3* and *GLI1/GLU3* loci ∼20 cM apart, indicating frequent recombination (6, 58) and the possibility to recombined *1AS^S263A^*and *1BS^S225E^* deletions with *GLI-A3* and *GLI-B3* alleles with reduced immunogenicity described below.

#### 3.6.1. GLI-A3

In Summit *GLI-A3*, the ω1,2-gliadins are pseudogenes with negligible expression in developing grains (Table S2) and encode proteins with early stop codons (Q91* and Q79*, Table S10) that eliminate ∼75% of the protein. These proteins are most likely not functional, so this allele was designated *gli-A3^null^*. Because the *1AS^S263A^* deletion is in the Summit background, it is already in phase with the *gli-A3^null^* locus, resulting in a CeD-safe combined region designated *1AS^null^*.

To facilitate the introgression of the combined *1AS^null^*region, we complemented the markers for *1AS^S263A^* (Fig. 4C-D) with two additional CAPS markers for *gli-A3^null^*. These markers use a single pair of primers (1AgliOAF1/OAR1, Table S6) and two different restriction enzymes. *Nco*I digestion distinguishes *gli-A3^nul^* alleles in Summit and Fielder (434 bp) from functional alleles in CS, RIL143, and Pegaso (248 + 186 bp; Fig. S8A), whereas *AlwNI* digestion differentiates Kronos (295 + 139 bp) from the other tested lines (434 bp; Fig. S8B). Combining the *gli-A3^null^* and *1AS^S263^* markers, we introgressed the *1AS^null^* region into Kronos by four backcrosses.

#### 3.6.2. GLI-B3/GLU-B2

To replace the Summit *GLI-B3* allele encoding proteins with 9 immunogenic epitopes, we identified two alternative *GLI-B3* alleles with no functional ω1,2-gliadins. The first, from tetraploid cv. Kronos, contains two ω1,2-gliadin pseudogenes and a linked functional LMW-GS gene (*GLU-B2* (6)) encoding a protein without known CeD immunogenic epitopes (Table S10). This locus was designated *gli-B3^KRO^*. A CAPS marker (GliOBF13/R13 with *Afe*I digestion, Table S6) differentiates *gli-B3^KRO^* (680 bp) from other *GLI-B3* alleles in hexaploid wheat (526 + 154 bp, Fig. S8C). Using these markers, we identified a BC_4_F_2_ plant homozygous for both *gli-B3^KRO^*and *1BS^S225E^* during backcrossing into Kronos, designated *1BS^S225E/Glu-B2^*. This recombinant 1BS carries no gliadins but retains two genes encoding functional LMW-GS lacking known CeD immunogenic epitopes (Summit *GLU-B3.4* and Kronos *GLU-B2*), so it was selected for the breeding objective. Although currently in tetraploid wheat, this recombined region is being introgressed into hexaploid wheat breeding lines.

Because the Kronos GLU-B2 LMW-GS in the recombined *1BS^S225E/Glu-B2^*may still harbor unidentified epitopes, we searched for *GLI-B3* alleles lacking all functional prolamins (*gli-B3^null^*) for the CeD-safe objective. We identified such an allele in CDC-Stanley (59), containing three ω1,2-gliadin pseudogenes with early premature stop codons and one LMW-GS pseudogene disrupted by repetitive elements (Table S10). We developed a CAPS marker (GliOBF14/R14 with *Nsi*I digestion, Table S6) to distinguish *GLI3* alleles of CDC-Stanely (461 bp) and Summit (335 + 140, Fig. S8D). Following CRISPR editing of the residual LMW-GS linked to *1BS^S225E^*, we plan to combine the edited *1BS^S225E^*with the *gli-B3^null^* to generate a *1BS^null^* region lacking prolamin genes. In summary, we completed the development of 1AS (1AS^null^) and 1DS (1DS^S204F^) chromosome arms lacking all prolamin genes and identified the components for a prolamin-free 1BS^null^ line.

### 3.7. RP-UPLC, A-PAGE, and immunoblot characterization *GLI1/GLU3/GLI3* prolamins

#### 3.7.1 RP-UPLC and A-PAGE

The RP-UPLC analysis of the gliadin fraction revealed multiple missing peaks in the deletion lines. The 1AS^S263A^ line lacked two γ-gliadin peaks but showed no changes in the ω1,2-region (Fig. 7A), consistent with the deletion of three γ-gliadin genes in *1AS^S263A^*(Table S9) and the absence of functional ω1,2-gliadin genes on Summit 1AS (Table S2). The two 1AS ω5-gliadins are weakly transcribed relative to its orthologs in other genomes (Table S2), correlating with the slight reduction in ω5-gliadins abundance (9.9% in 1AS^S263A^ vs. 12.5% in Summit WT, Fig. S6, Table S8) and the reduced intensity of one of the RP-UPLC peaks in the ω5-region (Fig. 7A).

**Fig. 7.**
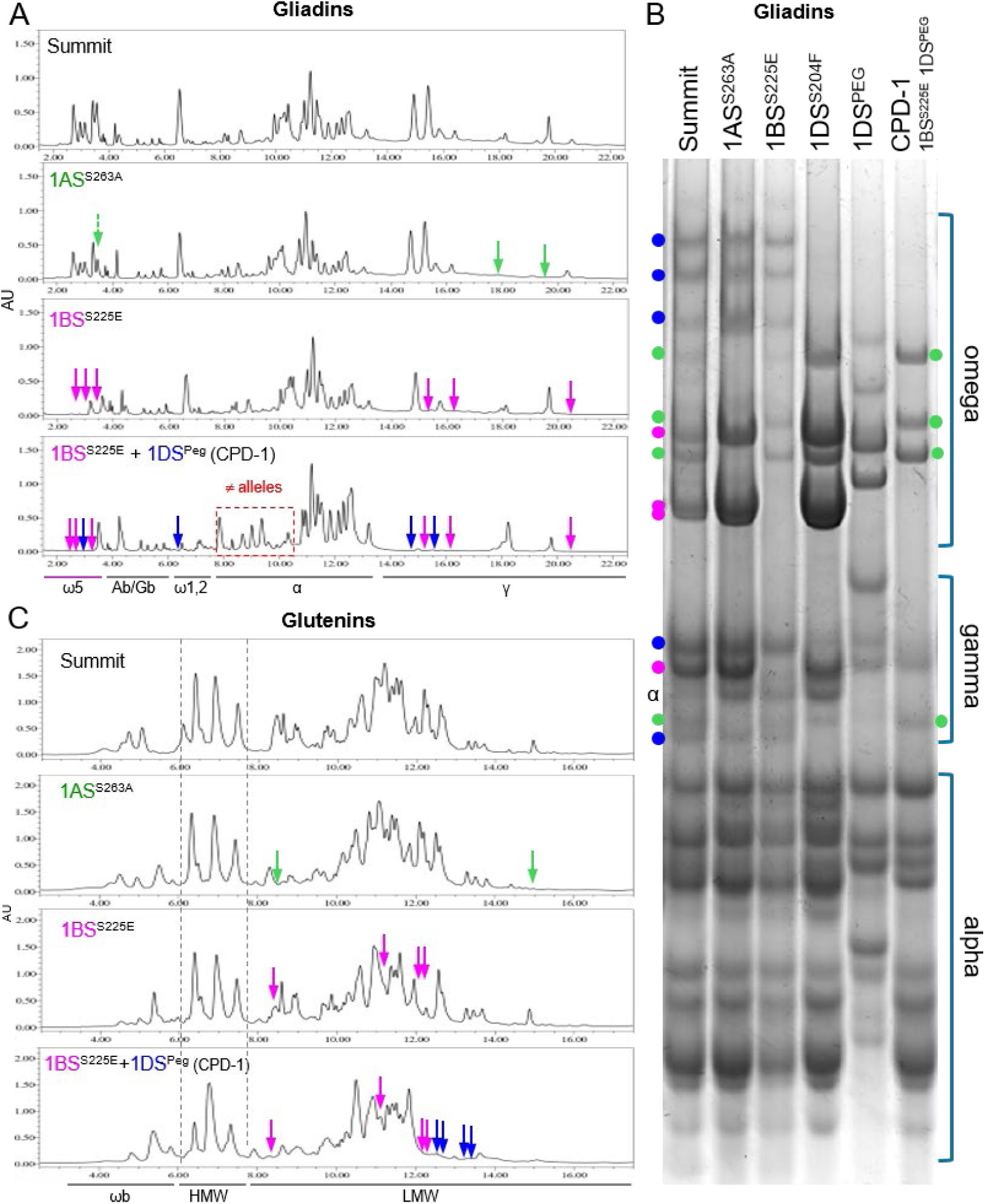
RP-UPLC and A-PAGE analyses for Summit WT and deletions including different *GLI1/GLU3* loci. Green arrows or dots indicate peaks or bands from the A genome, pink from the B genome and blue from the D genome (dotted arrows indicate RP-UPLC changes in relative intensity) (**A**) RP-UPLC analysis of lines lacking γ-gliadins and ω-gliadins. ω5= omega5-gliadins, Ab/Gb= albumins and globulins, α= alpha-gliadins, and γ= gamma-gliadins. (**B**) A-PAGE analysis of gliadins in the same lines. (**C**) RP-UPLC analysis of lines lacking LMW-GS. The 1DS^PEG^ line lacks some LMW-GS from *GLU-D3* but also includes other duplicated/polymorphic LMW-GS. HMW= High molecular weight glutenins, LMW= low molecular weight glutenins, and ωb= omega bound gliadins. The x-axis represents elution times (min), while the y-axis represents the intensity of the signal in absorbance unit (AU).

The A-PAGE showed loss of three ω- and one γ- bands in the 1AS^S263A^ line (Fig. 7B).

The 1BS^S225E^ line lacked three RP-UPLC peaks in both γ- and ω5-gliadin regions, which correspond to the four γ-gliadins and three ω5-gliadin genes present within the deleted region (Fig. 7A, Table S9), and the three ω- and one γ-gliadin bands missing in the 1BS^S225E^ A-PAGE profile (Fig. 7B). Because the only functional ω1,2-gliadin gene on 1BS lies outside the deletion (Table S9), no changes were observed in RP-UPLC ω1,2-gliadin region in 1BS^S225E^ (Fig. 7A).

Summit 1DS carries four highly transcribed ω1,2-gliadins (Table S2), corresponding to the major peak in the RP-UPLC ω1,2-gliadin region (Fig. 7A) and the three A-PAGE ω-gliadin bands (Fig. 7B) absent in 1DS^PEG^. This deletion also eliminated two γ-gliadin and one ω5-gliadin peaks in the RP-UPLC profile (Fig. 7A), and two γ-gliadin bands in the A-PAGE profile (Fig. 7B), corresponding to gliadin genes present within the deleted region (Table S9).

Line CPD-1 (*1BS^S225E^ 1DS^PEG^*) exhibited marked reductions relative to Summit in the RP-UPLC proportion of ω1,2-(12.5 to 2.8%), ω5-(6.2 to 3.4%), and γ-gliadins (33.4 to 9.1%, Fig. S6, Table S8). The A-PAGE profile of CPD-1 suggested that most remaining ω- and γ-gliadins bands originated from the intact *GLI-A1* locus (Fig. 7C). Overall, the single and combined deletions accounted for most ω- and γ-gliadin RP-UPLC peaks and A-PAGE bands.

In the glutenin fraction, the RP-UPLC identified two peaks missing in line 1AS^S263A^ and four in line 1BS^S225E^ (Fig. 7B), consistent with deletions of three and four LMW-GS genes, respectively (Fig. 7C and Table S9). Four additional peaks absent in CPD-1 relative to 1BS^S225E^ were likely associated with the *1DS^PEG^* deletion. Because this line still segregates for RIL143, Pegaso, and Summit genomes, the identities of deleted versus polymorphic LMW-GS remain unresolved.

Additional RP-UPLC peaks in CPD-1 likely originated from the duplicated *GLU-D3* region.

#### 3.7.2. Immunoblotting

To further characterize the ω-gliadins, immunoblots were performed using published ω5- and ω1,2-gliadin antibodies (Fig. 8). Profiles of these two antibodies in Summit and 1AS^S263A^ were highly similar (Fig. 8, 1^st^ row), consistent with the absence of functional ω1,2-gliadin genes and relatively low transcript levels of ω5-gliadin genes in Summit 1AS relative to the other homeologs (Table S2).

**Fig. 8.**
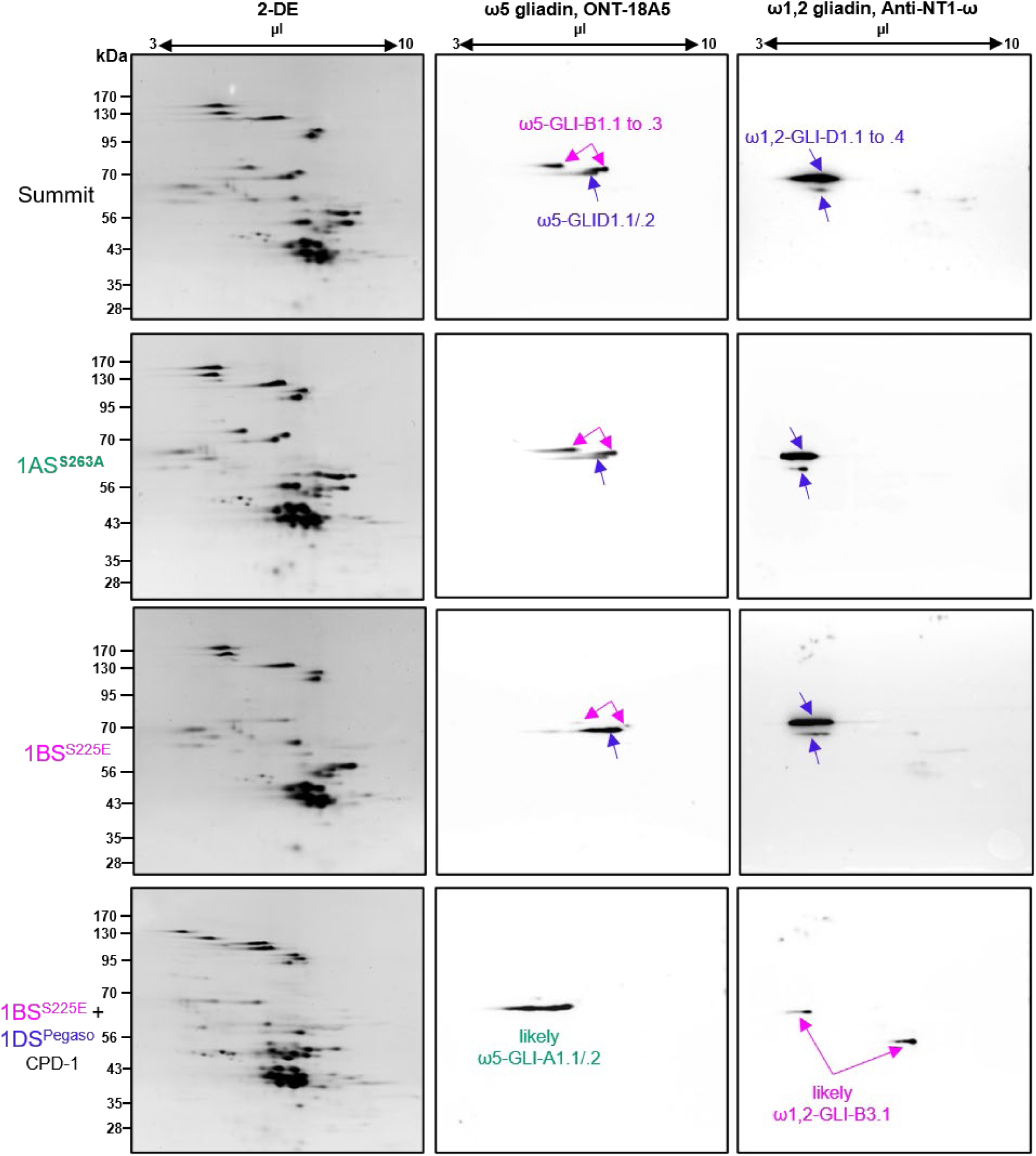
Immunoblots using ω5 (ONT 18A5) and ω1,2 (Anti-NT1-ω) antibodies. Left column: gliadin proteins separated by two-dimensional gel electrophoresis (2-DE). Central column: Protein spots reactive to ω5 gliadin monoclonal antibody (ONT 18A5, central column) were identified as 1B and 1D ω5-gliadins. Right column: Protein spots reactive to Anti-NT1-ω (right column) were identified as ω1,2-gliadins. Proteins encoded by genes in the A genome are indicated in green from the B genome in pink and from the D genome in blue. Proposed protein names are from Summit Table S2.

The Summit *GLI-B1* locus contains three highly expressed ω5-gliadin genes (Table S2), likely corresponding to the ω5 spots absent in the 1BS^S225E^ immunoblot (Fig. 8, row 3). Since Summit *GLI-B1* lacks functional ω1,2-gliadin genes (Table S2), no changes were expected for the 1BS^S225E^ proteins hybridized with the ω1,2 antibody.

Finally, most of the ω1,2- and ω5-gliadin proteins detected in Summit were absent in line CPD-1 (*1BS^S225E^ 1DS^PEG^*), whereas residual proteins showed compensatory increases in intensity (Fig. 8). Residual ω1,2-gliadin spots in CPD-1 most likely correspond to the functional ω1,2-GLI-B3.1 protein encoded by a gene outside the *1BS^S225E^* deletion (Fig. 8, Table S9), whereas residual ω5-gliadin spots probably derive from the two functional ω5-gliadins in 1AS, which are present in CPD-1 (Table S2 and S8). This interpretation is supported by the A-PAGE profiles (Fig 7C).

## 4. Discussion

A major challenge for reducing or eliminating wheat CeD immunogenic epitopes is the large number of prolamin genes encoding proteins with these epitopes (Table S2). However, because these genes are clustered in 11 regions (considering tightly linked *GLI1/GLU3* and *GLI3/GLU-B2* as single regions), the problem can be partitioned into smaller subprojects focused on generating individual CeD-safe regions that breeders can combine using molecular markers.

Since the *GLI1* and *GLI3* loci on chromosomes 1A and 1B are separated by only 10–20 Mb, the number of target regions can be reduced from 11 to 9 by selecting CeD-safe alleles for these two linked loci simultaneously with molecular markers. This strategy will enable the evaluation of individual prolamin loci and the development of optimal loci combinations that minimize immunotoxicity while preserving desirable end-use quality traits.

### 4.1. *GLI1/GLU3* and *GLI3/GLU2* linked loci

For chromosome arm 1AS, we identified a small deletion eliminating all functional prolamin genes from *GLI-A1/GLU-A3* (*1AS^S263A^*), confirmed its linkage to *gli-A3^null^*, and developed molecular markers for simultaneous selection of both loci. This prolamin-free *1AS^null^*region is ready to use for the development of CeD-safe genetic stocks. However, its use in breeding may be limited by the known negative effect of natural *Glu-A3e* alleles lacking functional LMW-GS on gluten strength and BMQ (60, 61). It would be important to investigate if these negative effects can be offset by the additional LMW-GS present in *1DS^PEG^* and/or by compensatory increases in glutenin expression associated with the elimination of highly expressed gliadins.

For chromosome arm 1BS, we identified a small *GLI-B1/GLU-B3* deletion (*1BS^S225E^*) lacking gliadins but retaining one LMW-GS gene outside the deletion (*GLU-B3.4*). This deletion was combined with the *Gli-B3^KRO^* allele, which also lacks gliadins and carries a functional LMW-GS (*GLU-B2*). The two retained LMW-GS in *1BS^S225E/Glu-B2^* lack known CeD immunogenic epitopes (54, 55), making this combined locus promising for the development of CeD-safe wheats. However, because the presence of unidentified immunogenic peptides cannot yet be excluded, we are editing the two genes outside the *1BS^S225E^* deletion and, in parallel, combining *1BS^S225E^*with the prolamin-free *gli-B3^null^* allele from CDC-Stanley.

For chromosome arm 1DS, we developed separate lines for the different objectives. For the CeD-safe objective, we selected the *1DS^S204F^*deletion that removes all known prolamins from 1DS. For breeding applications, we selected the *1DS^PEG^* chromosome that lacks all gliadins and carried duplicated LMW-GS. Although the complete set of LMW-GS sequences in *1DS^PEG^* is currently unavailable, orthologous genes in Summit and Fielder contain few or no CeD epitopes and lack immunodominant epitopes. This chromosome may therefore be useful for the development of breeding lines with reduced immunogenicity. However, recurrent seed purifications will be required to maintain this line because cytogenetic instability can cause secondary LMW-GS deletions. This instability may also facilitate selection of a *1DS^PEG^* chromosome lacking all LMW-GS genes for CeD-safe wheat development. Additional work will be required to generate a more stable 1DS stock lacking gliadins and immunogenic LMW-GS.

An unresolved question is whether combining the *1AS^S263A^*, *1BS^S225E^* and *1DS^S204F^* deletions will produce viable and fertile plants, since the deleted regions share null alleles for multiple non-prolamin genes. If the triple mutant proves inviable, more precise editing of gliadin genes for at least one *GLI1/GLU3* locus will be necessary. A recently published Fielder line with most of the ω- and multiple γ-gliadins edited by CRISPR (28) may provide a useful starting point.

### 4.2. *GLI2* loci

A previous study using γ-ray radiation mutants for *GLI-A2, GLI-B2*, and *GLI-D2* in RIL143 failed to combine all three alleles (12). This limitation was overcome here by combining *6DS^RIL143^* with smaller complementary *6AS^S288B^* and *6BS^S094D^* deletions in Summit. The resulting triple mutant was fertile and showed no obvious morphological defects, demonstrating the feasibility of this strategy. We retained the *6DS^RIL143^* deletion because in replicated field trials it positively affected BMQ and showed no negative impact on grain yield, either alone or combined with *6AS^RIL143^* (12). Combined deletions of the 6DS and 6AS α-gliadins remove 115 immunogenic epitopes (31% of all immunogenic epitopes, supporting the use of α-gliadin deletions as a primary target for reducing CeD immunogenicity in wheat breeding programs.

Line CPD-3 lacks 33 of the 35 α-gliadins present in Summit but retains two α-gliadins outside the *6BS^S094D^* deletion with two CeD immunogenic epitopes. Because neither epitope is immunodominant, this line may still be useful for breeding applications. However, for the CeD-safe wheat, we have initiated CRISPR experiments targeting these remaining α-gliadins.

### 4.3. *GLU1* loci

The *GLU1* loci are simpler to manipulate than other prolamin loci because each contains only two linked genes. We generated a triple mutant line combining *glu-A1^null^*, *glu-B1^K1280^*, and *1DL*^S059D^ deletion with no functional HMW-GS (*glu-A1^null^*, Fig. 1 and S3). This line was fertile, displayed no obvious morphological defects, and is suitable for CeD-safe wheat development. However, elimination of all functional *GLU1* genes is expected to substantially reduce BMQ, limiting its breeding utility. Potential strategies to reduce HMW-GS immunogenicity while maintaining gluten strength include prime editing (62) of *GLU1* immunogenic epitopes or replacement of the endogenous HMW-GS with synthetic *GLU1* genes with edited CeD-toxic epitopes. Both approaches are currently under investigation in our laboratory.

In summary, this study demonstrates that the complex problem of reducing CeD immunogenicity in wheat can be divided into smaller, tractable subprojects that can be precisely tackled with modern genomic technologies. This modular approach allows multiple laboratories to work in parallel to develop and evaluate new CeD-safe loci or optimize the ones described in this study and adapt them to specific research or breeding objectives. We are making the genetic stocks developed in this study publicly available to accelerate the development of CeD-safe research stocks and reduced immunogenicity breeding lines.

## Acknowledgements

We thank Gabriela Grigorean at the UC Davis Proteomics Center for her help with the proteomics analyses and Matthew Milner (USDA-ARS Western Regional Research Center) for reviewing the manuscript for readability.

## Data Availability

Raw data and statistical analysis are included in the Supplementary Data. The Summit genome reads and the RNA-seq data for Summit developing grains have been deposited in NCBI under BioProject number PRJNA1438777 (https://www.ncbi.nlm.nih.gov/sra/PRJNA1438777). The Summit genome assembly is available on NCBI (Genome JBXEOT000000000) and annotations are available at: https://zenodo.org/records/19598726. The genome can be visualized at https://graingenes.org/GG3/genome_browser. We have deposited 6DS^RIL143^ (PI 704908), Pegaso (PI 709644), Kronos *glu1^null^* (PI 692252) in GRIN-Global. The other lines will be deposited as soon as the USDA-ARS germplasm release review is completed.

## Author Contributions

MGR contributed to the experimental part, data analyses, and wrote the initial draft. DLC developed the Summit deletion lines and performed the RNA-seq and initial characterization of the deletion lines. WZ and XZ developed all the deletion combinations and generated all the molecular markers. MHGL characterized the Pegaso 1DS chromosome and provided A-PAGE and SDS-PAGE analyses. JZ, CC and GB provided bioinformatics and statistical analyses. SK and JYL performed the RP-UPLC and immunoblot assays. UY and EA sequenced, assembled and annotated the Summit genome. All authors contributed to the revision of the manuscript. JD proposed and coordinated the project, obtained funding, provided statistical analyses, supervised MGR, and wrote the final version of the paper.

## List of supplemental materials

### Supplemental Tables

Table S1 Summit genome sequencing statistics

Table S2 Summit prolamins

Table S3 RNA-seq statistics

Table S4 Exome capture sequencing statistics

Table S5 Borders 1DL deletion

Table S6 Primers used in this study

Table S7 Borders 6AS and 6BS deletions

Table S8 Quantification of RP-UPLC peaks

Table S9 Borders homeologous group 1

Table S10 Sequences of genes in *GLI3* and *GLU-B2* loci

Table S11 Prolamin proteins reference for proteomics analysis

Table S12 Spectral counts of peptides diagnostic for ω1,2-gliadins at the *GLI3* locus

### Supplemental Figures

Figure S1 Genome assembly alignments

Figure S2 Grain expression profiles

Figure S3 Generation of glu1^null^

Figure S4 6AS deletion in Pegaso

Figure S5 Generation of combined *gli2* deletions CPD-2 and CPD-3

Figure S6 RP-UPLC Quantification

Figure S7 Cytogenetic instability of 1DS^Peg^

Figure S8 Comparisons among ω1,2-gliadins encoded be genes in *GLI1* & *GLI3* loci

Figure S9 Molecular markers for the *GLI-A3* and *GLI-B3* loci

