## Supplementary figures for "A genetic toolkit to reduce wheat immunogenicity and incidence of celiac disease"

**Figure S1.** Alignment of the Summit and Kariega (https://doi.org/10.1038/s41588-022-01022-1) genome assemblies.

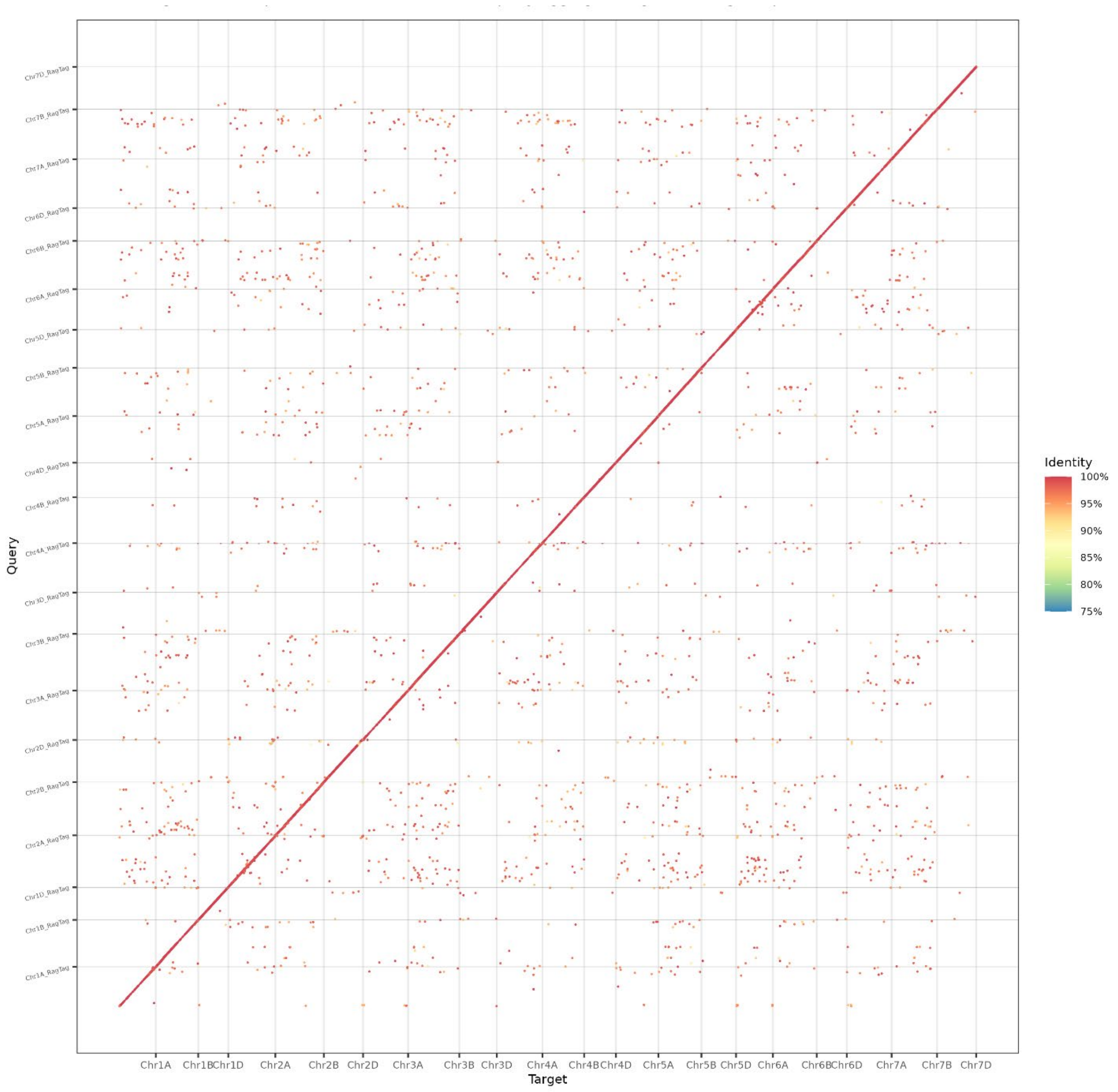

**Figure S2.** Expression profiles of different prolamin genes in Summit developing grains at 14, 21, 28 and 35 days post anthesis (dpa). (**A**) Expression graph showing the dynamic changes in expression of the different classes of prolamins during grain development in Summit. (**B**) Principal component analysis of the different prolamin classes and different genomes. The first PC explains 86.9% of the variation and reflects mainly differences in transcript levels, whereas the second PC explains 9.6% of the variation and is mainly driven by differences in the time of peak expression.

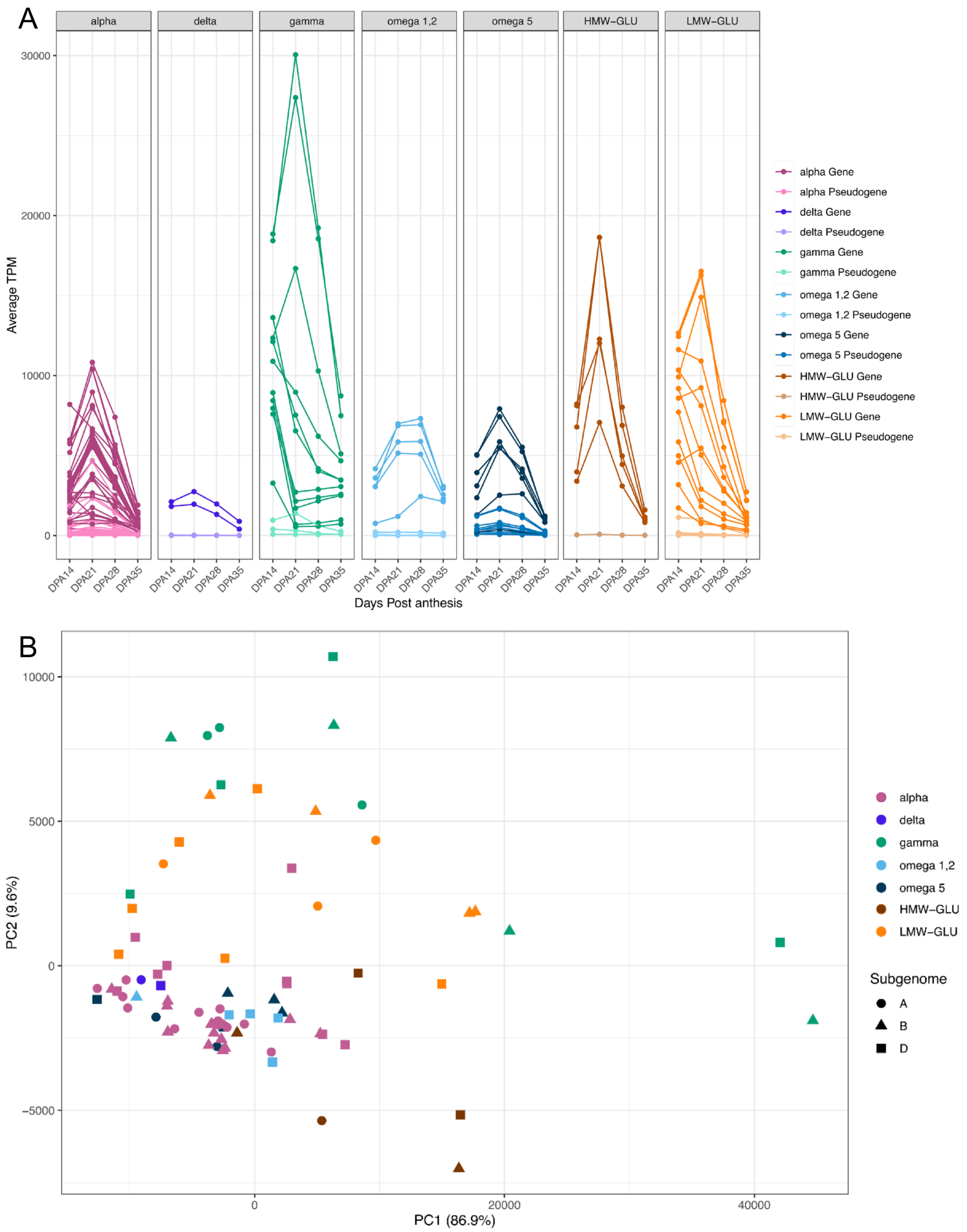

**Figure S3.** Generation of a line without functional HMW-GS. Summit line 1DL$^{S059D}$ (1DL deletion encompassing the *GLU-D1* locus) was crossed with Kronos EMS mutant line glu-B1$^{K1280}$ (premature stop codons in both *Bx* and *By* genes). Kronos has a natural non-functional *GLU-A1* locus, so both *glu-A1$^{null}$* and *glu-B1$^{K1280}$* alleles from this line were transferred to hexaploid wheat. The pentaploid $F_1$ was backcrossed four times to Summit 1DL$^{S059D}$, selecting for heterozygous *glu-A1$^{null}$* and the *glu-B1$^{K1280}$* alleles in each generation using molecular markers (Table S6). $BC_4$ heterozygous plants were self-pollinated and the triple homozygous $BC_4F_2$ line, designated as glu1$^{null}$ was selected using the same molecular markers.

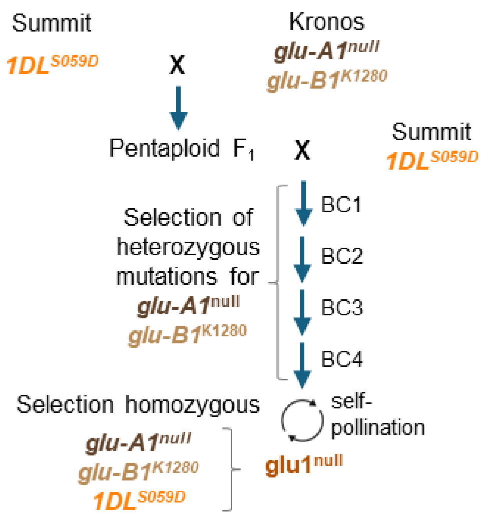

**Figure S4.** Deletion in Pegaso chromosome 6AS encompassing the *GLI-A2* locus. The 6AS deletion in Pegaso (*6AS*$^{PEG}$), transferred from the US cultivar Reader (Camerlengo et al. 2017, BMC Plant Biol, 17:248), is 41.56-41.68 Mb long (Table S7). Pegaso exome capture reads were mapped to the Summit genome and compared with a Summit exome capture mapped to the same genome. The deletion graph below presents the ratio between the normalized reads per gene in Pegaso / normalized reads per gene in Summit using a sliding window of 10 genes to obtain smoother curves.

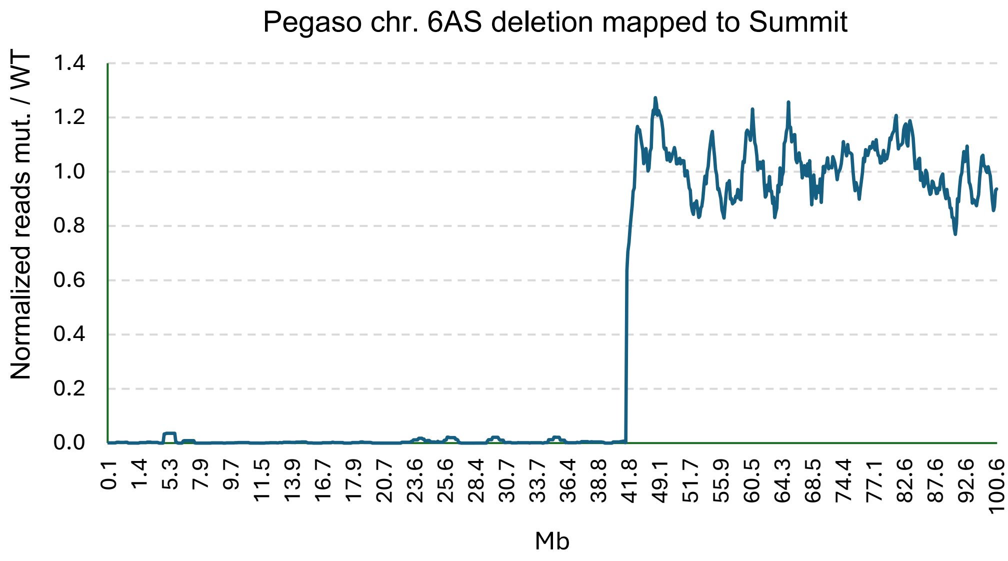

**Figure S5.** Combination of deletions for HMW-GS and α-gliadins. We first crossed glu1$^{null}$ with 6DS$^{RIL143}$ and backcrossed the $F_1$ four times to Summit glu1$^{null}$ to recover the Summit genetic background. We selected a $BC_4F_2$ plant homozygous for *glu1*$^{null}$ and *6DS*$^{RIL143}$ and crossed it with glu1$^{null}$ 6AS$^{S288B}$. From the $F_2$ progeny of this cross we selected a plant homozygous for *6AS*$^{S288B}$, *6DS*$^{RIL143}$, and the three non-functional HMW-GS alleles, which was designated as CPD-2. Finally, we crossed CPD-2 with 6BS$^{S094D}$, and in a large $F_2$ progeny selected a line homozygous for the three *GLI2* deletions plus the *glu-B1*$^{K1280}$ and *1DL*$^{S059D}$ alleles. The functional *GLU-A1x* gene is still present in this line designated as CPD-3.

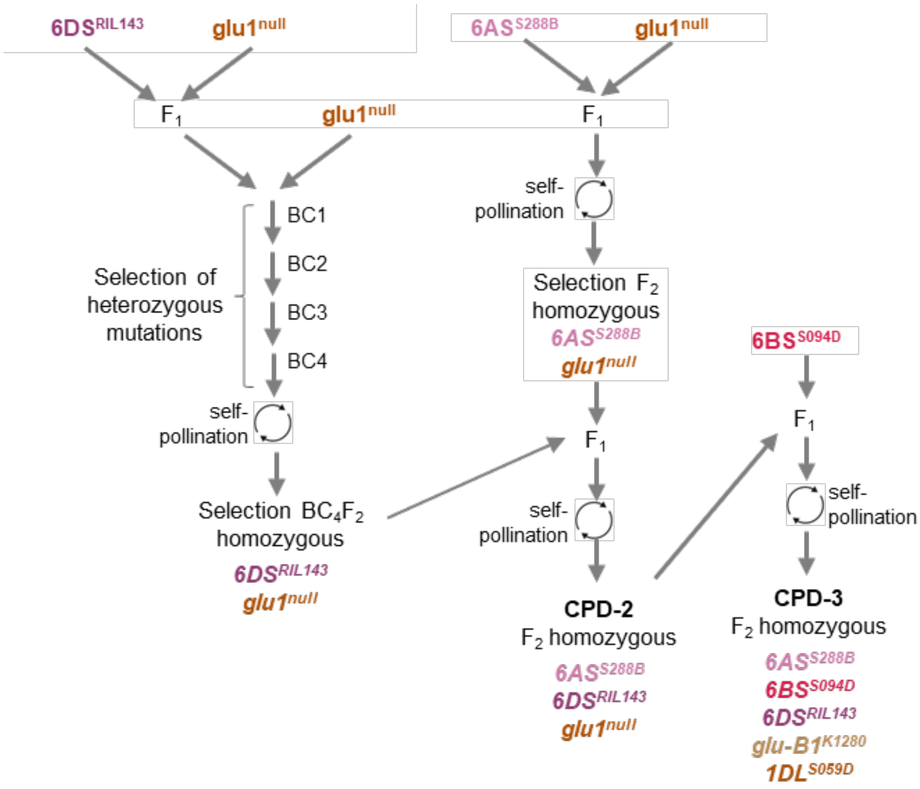

**Figure S6.** Quantification of the RP-UPLC [%] fractions in Summit and deletion lines. (**A-B**) Relative amount of ω5-, ω1,2-, α- and γ gliadins. (**C-D**) Relative amounts of LMW-GS and HMW-GS. (**A** and **C**) Relative percentage within each line. (**B** and **D**) Relative percentages normalized to Summit. Data presented as mean value of 3 replications. Raw data available in Table S8. Line descriptions:

1. SUM: Summit wt.
2. glu1$^{null}$: no functional HMW-GS.
3. CPD-2: *6AS*$^{S288B}$ (*GLI-A2*), *6DS*$^{RIL143}$ (*GLI-D2*), *glu1*$^{null}$ (non-functional *glu-A1, B1,* & *D1*).
4. 6BS$^{S094D}$: *GLI-B2* deletion (2 α-gliadins outside the deletion).
5. 1AS$^{S263A}$: *GLI-A1* deletion linked to no-functional *GLU-A3* locus.
6. 1BS$^{S255E}$: *GLI-B1* deletion linked to *GLU-B3* (one LMW-GS) and *GLI-B3* (one ω1,2-gliadin).
7. 1DS$^{PEG}$: *GLI-D1* deletion and *GLU-D3* duplication.
8. CPD-1: *1BS*$^{S255E}$ and *1DS*$^{PEG}$ combined deletions.

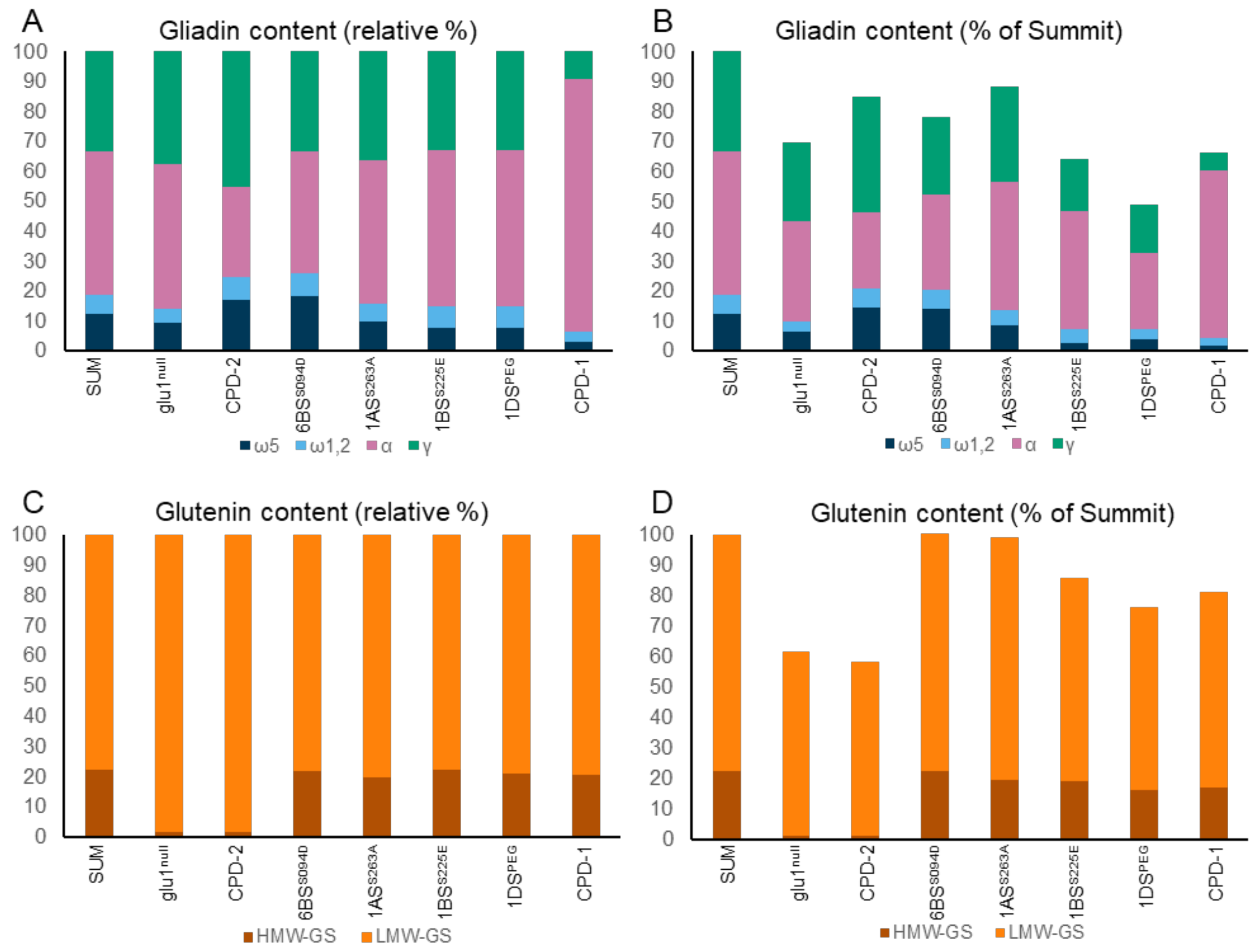

**Figure S7. Cytogenetic instability of the 1DS$^{PEG}$ chromosome.** (**A-B**) Marker GligDF5/DR5del1D (Table S6) for the *GLI-D1* deletion confirmed that 1DS$^{PEG}$ and all 10 plants from head-rows 103 (**A**) and 108 (**B**) were homozygous for the deletion. Summit shows the expected 755 bp band and serves as positive control. (**C-D**) Marker 1Dglu3F3/R3-LMW-het (Table S6) amplified two duplicated LMW-GS that are separated after digestion with *Nco*I in line 1DS$^{PEG}$, and a single fragment in Summit. Among the 10 plants from head-row 103 (**C**), we detected two plants with a single fragment, and among the 10 plants from head-row 108 (**D**) we found four plants with no amplification. The other 80 plants showed amplification of both bands. (**E**) Model to explain the origin of these novel 1DS$^{PEG}$ chromosomes. A recombination event within inverted segments in two 1DS$^{PEG}$ chromosomes paired in inverted orientation results in a bridge-and-fragment. Random breaks in the dicentric bridge during meiosis can explain both the single band in (**C**) and the absent bands in (**D**). The two yellow arrows in the bridge represent independent breaking point in different individuals.

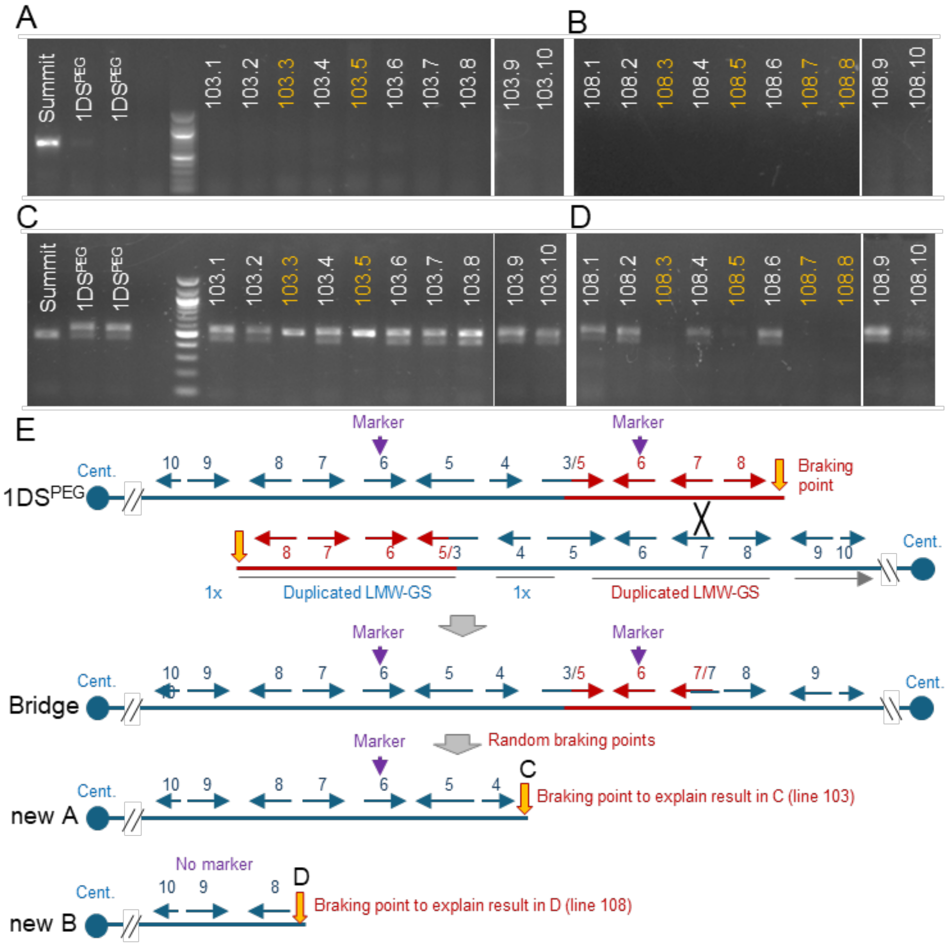

**Figure S8.** Comparisons among GLI1 and GLI3 ω1,2-gliadins. (**A**) Neighbor-Joining tree for ω1,2-gliadin proteins located in the *GLI1* and *GLI3* loci [1]. Branch lengths are proportional to the evolutionary distances used to infer the tree, and evolutionary distances are in number of amino acid substitutions per site. All ambiguous positions were removed for each sequence pair (pairwise deletion option). There were a total of 435 positions in the final dataset. Evolutionary analyses were conducted using MEGA11 [2]. (**B**) Alignment of the C-terminal regions of ω1.2-gliadins showing premature stop codons in GLI-A3 and GLI-B3 proteins. The ω1.2-gliadin designations are based Chinese Spring (CS) curated prolamin annotations and independent sequential number in Fielder and Summit (same numbers do not indicate orthologous genes). Kronos (KRN) numbers are based on the CS prolamin annotation. The functional GLI-B3 ω1.2-gliadin proteins are identical in CS (ω1,2-GLI-B3.9), Fielder (ω1,2-GLI-B3.11), and Summit (ω1,2-GLI-B3.1), but the KRN ortholog is a pseudogene with a premature stop at protein position 250 (linked to a functional LMW-GS).

A

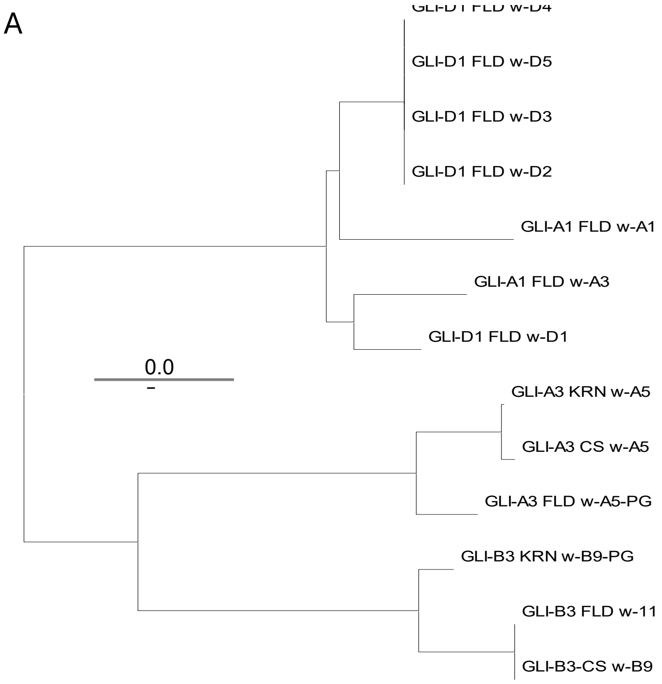

B

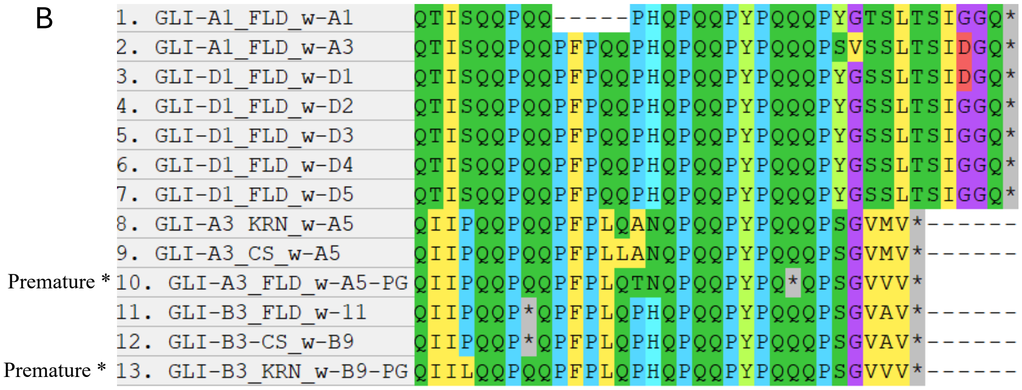

**Figure S9. Molecular markers for the *GLI-A3* and *GLI-B3* loci**. (**A-B**) CAPS markers for the *GLI-A3* locus PCR amplified with primers 1AgliOAF1/OAR1 (Table S6) and digested with two different restriction enzymes. (**A**) Digestion with *Nco*I differentiates the *gli-A3*$^{null}$ allele present in Summit and Fielder form the functional *GLI-A3* alleles in CS, RIL143 and Pegaso, but it does not differentiate Summit from Kronos. (**B**) Digestion with *AlwN*I differentiates Kronos from all other tested lines. This marker is useful to introgress the *gli-A3*$^{null}$ allele into Kronos. (**C**) CAPS marker for the *GLI-B3* locus amplified with primers GliOBF13 /R139 and digested with *Afe*I. This codominant marker differentiates Kronos from the hexaploid varieties and is useful to introgress the Kronos *GLI-B3* allele with a functional LMW-GS (no epitopes) and non-functional ω1.2-gliadins into hexaploid wheat. (**D**) CAPS codominant marker amplified with primers GliOBF14/R14 and digested with *Nsi*I. This marker is useful to introgress the *gli-B3*$^{null}$ allele from CDC-Stanely and to recombine this allele with the *GLI-B1/GLUB3* deletion in line 1BS$^{S225E}$.

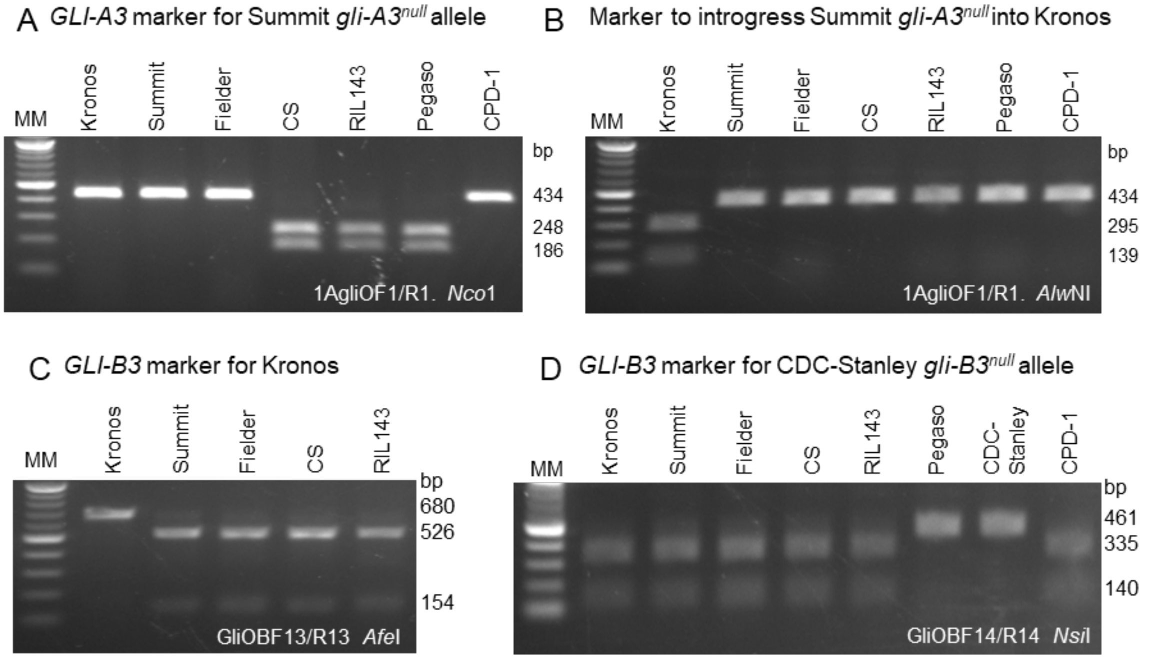
